# Dissecting the *cis*-regulatory syntax of transcription initiation with deep learning

**DOI:** 10.1101/2024.05.28.596138

**Authors:** Kelly Cochran, Melody Yin, Anika Mantripragada, Jacob Schreiber, Georgi K. Marinov, Sagar R. Shah, Haiyuan Yu, John T. Lis, Anshul Kundaje

## Abstract

Despite extensive characterization of mammalian Pol II transcription, the DNA sequence determinants of transcription initiation at a third of human promoters and most enhancers remain poorly understood. We trained and interpreted a neural network called ProCapNet that accurately models base-resolution initiation profiles from PRO-cap experiments using local DNA sequence. ProCapNet learns sequence motifs with distinct effects on initiation rates and TSS positioning and uncovers context-specific cryptic initiator elements intertwined within other TF motifs. ProCapNet annotates predictive motifs in nearly all actively transcribed regulatory elements across multiple cell-lines, revealing a shared *cis*-regulatory logic across promoters and enhancers and a highly epistatic sequence syntax of cooperative and competitive motif interactions. ProCapNet models of steady-state RAMPAGE profiles distill initiation signals on par with models trained directly on PRO-cap profiles. ProCapNet learns a largely cell-type-agnostic *cis*-regulatory code of initiation complementing sequence drivers of cell-type-specific chromatin state critical for accurate prediction of cell-type-specific transcription initiation.

## Introduction

Regulation of gene expression is central to development, cellular differentiation, homeostasis, and response to stimuli, while its dysregulation plays a causal role in disease. Transcription initiation is a pivotal process in gene regulation. The recruitment of the transcriptional machinery and the kinetics of transcription initiation are regulated by the coordinated activity of signaling inputs that direct combinatorial transcription factor (TF) occupancy and chromatin states at proximal and distal regulatory elements ^1,2^.

The core components of the transcriptional machinery in most metazoans, essential for transcribing protein-coding genes and most lincRNAs, include RNA Polymerase II and general transcription factors (GTFs), which form the preinitiation complex (PIC) ^3–21^. These protein complexes decode promoter sequences to precisely recruit and activate Pol II via interactions with promoter sequence elements, stabilizing interactions with other PIC proteins, and DNA unwinding at the transcription start site.

However, our understanding of promoter sequence elements that regulate transcription initiation is incomplete. Previously identified core promoter elements ^22^ include the TATA box ^23–25^, the initiator element (Inr) ^26–28^, the B recognition element (BRE) ^29,30^, and the “motif ten” element (MTE) ^31,32^. The contribution of other sequence-specific TF motifs in regulating initiation is largely unknown, with some notable exceptions such as NFY (CCAAT box) ^33,34^, SP1^35^, the ETS family members ^36^, and YY1^37–39^. Although some motif spacing constraints have been observed in promoters, such as the preferential positioning of the TATA box ∼30bp upstream of the Inr ^40,41^, the role of higher-order motif arrangements (syntax) in the regulation of transcription initiation is poorly understood. Furthermore, it is now established that tran-scription is not exclusive to gene promoters but is, in fact, also widespread at distal enhancers ^42–48^. However, it is unclear whether promoters and enhancers harbor distinct or shared initiation sequence codes.

Experimental efforts to functionally characterize initiation-relevant sequence elements and *cis*-regulatory logic have generally confirmed their importance, although on a limited scale. Saturated mutagenesis experiments on genomic sequences are difficult to scale genome-wide and generally measure only the first-order effects of individual base mutations ^49,50^, while massively parallel reporter assays measuring promoter activity remove the promoter from its endogenous chromatin and distal enhancer context ^51,52^; neither approach typically incorporates measurement of nascent transcription specifically.

To address these limitations, computational methods have been developed to characterize the sequence basis of initiation ^53,54^. The first wave of approaches searched for statistically overrepresented sequence features and enriched co-occurrence and spatial patterns of features in promoters ^31,41,55,56^. These observational enrichment methods are inherently limited in that 1) they are often not cell-context aware, and 2) noting the enrichment of a motif in promoters or near TSSs neither guarantees that the motif plays any role at all, nor describes what that role might be: does the motif’s presence influence the rate of PIC recruitment, the positioning of the PIC and thereby the choice of TSS, or both, and how?

Supervised machine learning methods have been proposed as an alternative approach, where computational models are trained to predict biochemical readouts of transcriptional regulation as a function of the underlying DNA sequence. Deep neural networks have proven to be particularly adept at simultaneously learning predictive sequence features and their higher-order syntax *de novo* from DNA sequence and accurately mapping the learned representations to various biochemical readouts, including TF binding ^57^, RNA splicing ^58^, and chromatin accessibility^59^. Model interpretation methods have been developed to infer the impact of individual sequence features, their syntax and other sequence variation specifically on the experimental read outs the models are trained on, thereby alleviating some of the above mentioned issues with enrichment methods ^57–59^.

Recently, large, multi-task, deep neural networks trained on steady-state RNA abundance from RNA-seq, CAGE, or RAMPAGE experiments have achieved impressive predictive performance ^60–64^. However, these models have provided limited insights into the *cis*-regulatory code of initiation partly due to challenges with scalable, robust interpretation of massive deep learning models, but also due to the sheer complexity of multiple layers of regulation between sequence and steady-state gene expression that these models attempt to learn *de novo* ^65,66^. While initiation determines the precise location of the transcription start site (TSS), the rate of eukaryotic transcription is predominantly determined by promoter-proximal pausing limiting re-initiation, rather than PIC recruitment rates ^67^, and post-elongation RNA processing steps (splicing, polyadenylation, etc.) can drastically impact both the products of transcription and their lifespans ^68–72^. Major differences between measurements of transcription made by steady-state assays vs. nascent transcription assays such as GRO-seq ^73^, PRO-seq/PRO-cap ^74^, CoPRO ^75^, and NET-seq ^76^ highlight how the composition of the steady-state transcriptome is determined by much more than the outcome of transcription initiation. Any attempts to model steady-state expression data, therefore, must necessarily model promoter-proximal pausing, RNA processing, and degradation in order to make accurate predictions. Recent attempts at this integrated approach have resulted in more accurate models that continue to pose challenges for robust and scalable interpretation ^63,64^. Scaling back to the intermediate goal of directly modeling transcription initiation with tractable, interpretable models is, therefore, a potentially promising approach. A recent study by Dudnyk *et al*. models base-resolution initiation profiles using a transparent, additive neural network architecture initialized with manually curated motif representations distilled from a larger, long-context model ^77^. While the rationale for this ad-hoc dual-model distillation approach is a purported trade-off between prediction performance and interpretability, it is unclear if these design choices actually enhance interpretation of the *cis*-regulatory sequence code of nascent transcription relative to well-established alternative approaches for model interpretation. Additionally, the study 1) models initiation profiles averaged over diverse cell types, thereby losing context specificity, 2) is limited to the analysis of promoters, and 3) solely focuses on the shape of initiation profiles, neglecting initiation rates and predictions of overall promoter strength.

Here, we adopt an alternative approach by developing ProCapNet, a compact, deep learning model coupled with a well-established and robust interpretation framework for deciphering the *cis*-regulatory sequence code of transcription initiation. ProCapNet accurately models base-resolution initiation profiles from cell-type-specific PRO-cap experiments, distinguishing between the rate of initiation and TSS positioning, using local DNA sequence. We interpret ProCapNet to derive a comprehensive sequence motif lexicon of transcription initiation that includes known and novel variants of core promoter motifs and other specific TF motifs, some of which preferentially impact initiation rates or TSS positioning. ProCapNet reveals cryptic Inr-like initiator codes intertwined within motifs of other TFs, which precisely position initiation events only in specific sequence contexts. ProCapNet identifies predictive motifs in nearly all actively transcribed regulatory elements, including a shared initiation sequence logic across promoters and enhancers with subtle syntactic differences in motif density, diversity and affinity, thereby providing a comprehensive genome-wide annotation of sequence drivers of initiation in 6 ENCODE cell-lines. ProCapNet also reveals extensive cooperative and competitive effects of motif combinations suggestive of a highly epistatic initiation sequence syntax. Training ProCapNet on steady-state gene expression profiles from CAGE and RAMPAGE assays enables *de novo* distillation of initiation activity that is remarkably concordant with predictions from models trained directly on PRO-cap profiles. Finally, we show that ProCapNet learns a largely cell-type-agnostic *cis*-regulatory code of initiation that complements sequence drivers of cell-type-specific chromatin accessibility. We identify chromatin state modalities critical for predicting the cell-type-specificity of transcription initiation.

## Results

### ProCapNet accurately predicts initiation activity and precise positioning of human TSSs from DNA sequence

ProCapNet is a convolutional neural network that models base-resolution transcription initiation profiles, measured by PRO-cap experiments, in 1 kb genomic segments as a function of local (2 kb) DNA sequence context (**Fig. 1A**) ^74^. ProCapNet’s architecture is based on the BPNet model previously developed to predict base-resolution TF binding profiles ^57^. We adapted BPNet to PRO-cap read-outs by increasing the number of parameters and modifying the loss function to account for the strand specificity of transcription initiation (see Methods). ProCapNet preserves a key modeling choice introduced in BPNet, to separately predict the total coverage and shape of base-resolution PRO-cap read count profiles over each genomic segment. The coverage of aggregated read-counts (in *log* scale) over each segment serves as a proxy for total initiation activity (rate). A multinomial probability distribution of read-counts over all bases across both strands of each segment is used to model the profile shape, which captures solely the strand-specific positioning of initiation events at putative TSSs. Explicit separation of total initiation activity from TSS positioning enables downstream interpretation of the contribution of learned sequence features to these two complementary properties of initiation profiles.

**Figure 1:**
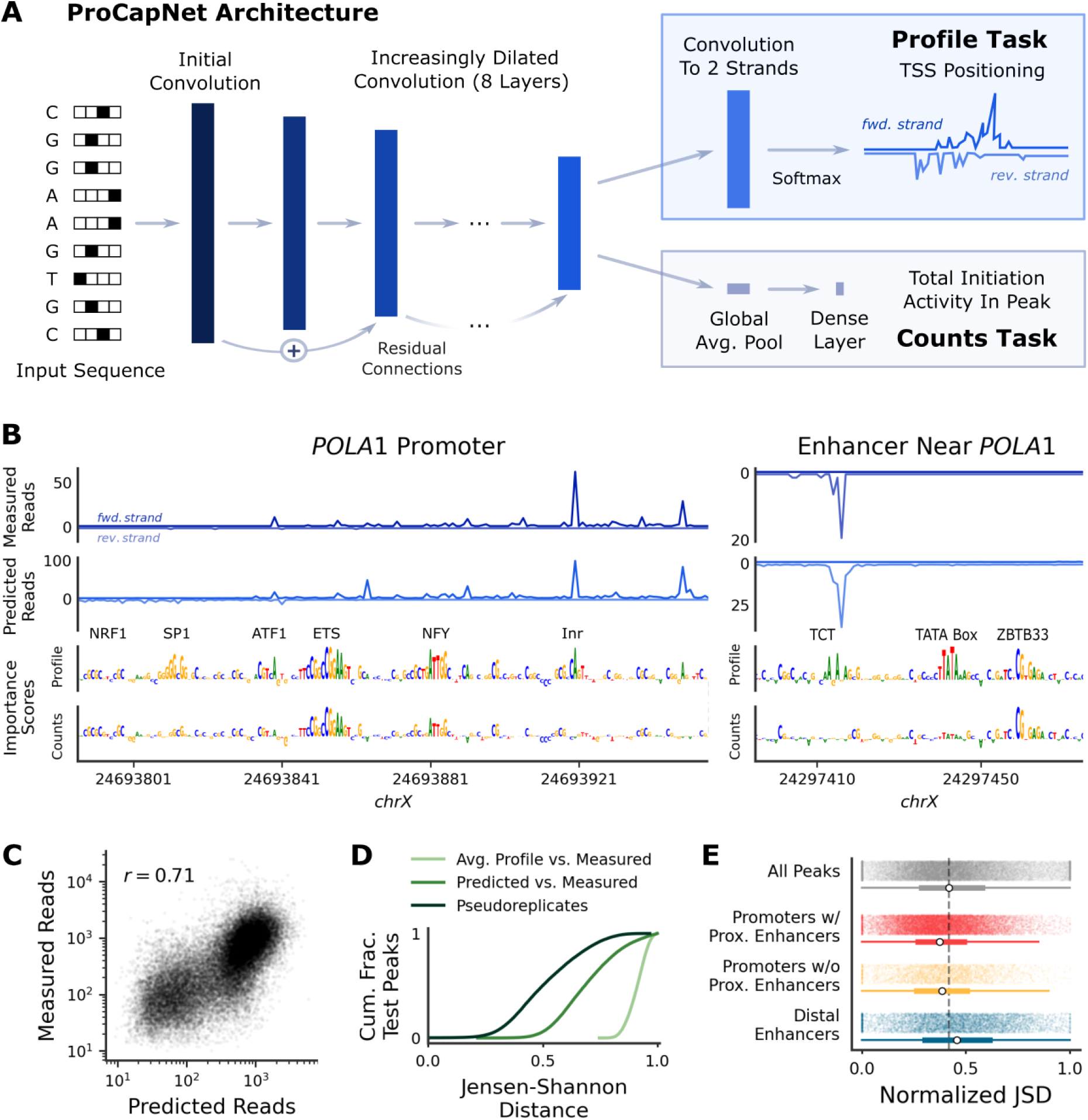
ProCapNet accurately predicts initiation rates and TSS positioning at base-pair resolution. (A) Design of ProCapNet, which is trained to predict base-resolution initiation events measured by PRO-cap. (B) Measured data, ProCapNet predictions, and ProCapNet contribution scores (for the profile and counts tasks) for two example PRO-cap peaks. Names of motifs highlighted by contribution scores are annotated. (C) Counts task performance on PRO-cap peaks from held out test chromosomes across 7-fold cross-validation. *r*, Pearson correlation. (D) Profile task performance on PRO-cap peaks from held out test chromosomes across 7-fold cross validation (higher JSD is worse). The *y*-axis shows the cumulative fraction of test peaks. (E) Profile task performance on held-out test peaks, stratified by overlap with ENCODE-annotated *cis*-regulatory elements. White dots are medians; the dashed line is the median for all test peaks. w/, with; w/o, without; Prox., proximal. JSD is normalized per-example to reduce confounding by coverage (see Methods; higher normalized JSD is worse).

We trained ProCapNet on genomic segments overlapping PRO-cap peaks and background regions in the human K562 cell-line, using a 7-fold cross-validation scheme. Since a vast majority (*>* 90%) of the PRO-cap peaks directly overlap or are within 500bp of DNase-seq peaks, we deliberately used accessible chromatin (DNase-seq peaks) lacking initiation as background regions to encourage the model to learn a *cis*-regulatory code of initiation decoupled from chromatin accessibility. We analyze the consequences of this choice later in the manuscript. Visual inspection of measured and predicted PRO-cap profiles at the *POLA1* promoter and a nearby enhancer highlighted strong correspondence (Pearson correlations *r* = 0.87 and 0.95 at the promoter and enhancer, respectively) of local maxima and strand specificity (**Fig. 1B**). Systematic evaluation of prediction performance across held-out PRO-cap peaks in cross-validation test sets corroborated the strong correlation (*r* = 0.72 *±* 0.025 across folds) between total measured and predicted coverage (*log* scale), relative to an upper-bound (*r* = 0.84) of correlation of measured coverage across the same peaks between replicate experiments (**Fig. 1C**). This correlation remained consistent (*r* = 0.72 *±* 0.015 across folds) across PRO-cap peaks and non-overlapping, accessible background regions, and the model’s predictions distinguished PRO-cap peaks from these background regions with an average precision score of 0.79 *±* 0.014 across folds. The gap between model performance and replicate concordance, also observed for BPNet TF binding models ^57^, is to be expected since ProCapNet, by design, only models the impact of local sequence context on PRO-cap readouts and hence cannot account for the potential effects of distal regulatory sequences and other non-sequence-based *cis* and *trans*-regulatory factors ^78^. ProCapNet’s base-resolution profile shape predictions in PRO-cap peaks were more similar (mean Jensen-Shannon Distance (JSD) = 0.69 *±* 0.004 across folds, lower JSD indicates higher similarity) to the measured profile shapes than a baseline prediction corresponding to the average PRO-cap profile across all peaks excluding those in the test set (mean JSD = 0.91; **Fig. 1D**). However, model performance was worse than measured profile shape concordance (mean JSD = 0.51) between pseudo-replicates of the same PRO-cap experiment. Together, these results indicate that ProCapNet can predict the magnitude and shape of PRO-cap profiles with substantial accuracy.

Next, we evaluated variability of model performance across subsets of PRO-cap peaks overlapping different classes of ENCODE candidate *cis*-regulatory elements (cCREs) ^79^. Profile prediction performance at PRO-cap peaks overlapping candidate distal enhancers was on par with peaks overlapping candidate promoters, and profile prediction performance at promoters was not dependent on the presence of a proximal enhancer nearby (**Fig. 1E**). The total coverage prediction performance (*r* = 0.48) at candidate promoters with proximal enhancers was higher than performance (*r* = 0.42) at candidate promoters without proximal enhancers, suggesting that ProCapNet’s coverage prediction performs best at promoters when a proximal enhancer is close enough to be included in the model’s input sequence (**Fig. S1A**). Total coverage performance across candidate distal enhancers was higher (*r* = 0.57) than performance across both classes of candidate promoters, suggesting that long-range regulatory interactions may have less influence on initiation at distal enhancers compared to promoters.

We also evaluated variability of model performance across subsets of PRO-cap peaks overlapping promoters of housekeeping genes, all protein-coding promoters, lncRNA promoters, promoters of ribosomal protein genes with TCT sequences, TATA-containing promoters, GENCODE annotations (i.e. in exons, in introns, or intergenic), and areas of high vs. low unique read mappability. We observed comparable profile shape prediction performance across most of these categories, with the exception of low-mappability sites which showed a minor deterioration of performance (**Fig. S1B-C**).

Finally, we evaluated the performance of ProCapNet accounting for two important attributes of transcription initiation events - directional imbalance and dispersion. Mammalian transcription is generally bidirectional, with substantial variability of directional imbalance of initiation across transcriptionally active loci ^47,73,80,81^. Initiation can also vary widely in terms of dispersion: at some loci, all or nearly all transcription initiates at a single base, while at other loci initiation can occur across a more dispersed window, up to 140 bp wide, along one strand ^22,48,82^. We quantified strand asymmetry of bidirectional initiation using the Orientation Index (OI), which is the fraction of reads on the majority strand ^83^. We quantified dispersion of initiation using an entropy-based measure called the Normalized Shape Index (NSI), such that higher values indicate more dispersion (modified from Hoskins *et al*. 2011^84^). Profile prediction performance of ProCapNet remained consistent across the spectrum of OI and NSI, with the exception of a small group of outliers where nearly all initiation occurred in one direction (OI *>* 0.9; **Fig. S1D**). Specifically, nearly half of the PRO-cap peaks with the most discordant profile predictions (normalized JSD *>* 0.9, bottom 6%, 1748 of 30,534 peaks; see Methods for JSD normalization details) were exclusively unidirectional (OI = 1). Compared to all other PRO-cap peaks, these outlier peaks had on average an order of magnitude fewer reads, were half as likely to overlap any annotated TSS, were twice as likely to be in intergenic regions or gene bodies, and particularly in exons and UTRs, and rarely overlapped annotated CREs (**Fig. 1E**; **Fig. S1E**). These trends suggest that at least some of these sites may have artefactual origins, such as from RNAs undergoing internal re-capping. This phenomenon has been previously reported in CAGE data ^85^; however, the extent of internal re-capping in PRO-cap-like assays is lesser than in CAGE due to experimental protocol differences, such as PRO-cap’s lack of size selection ^86,87^.

In summary, ProCapNet learns a unified, generalizable mapping of DNA sequence to initiation profiles with largely consistent predictive performance across diverse strata of genomic loci.

### ProCapNet reveals a comprehensive lexicon of sequence motifs predictive of transcription initiation rates and TSS positioning

Next, we used well-established model interpretation methods to decipher the *cis*-regulatory sequence features learned by ProCapNet ^57^. Using the DeepSHAP feature attribution method, we estimated the contribution of each base in the sequence of individual PRO-cap peaks to the total predicted coverage and profile shapes separately ^88,89^. DeepSHAP contribution scores at the *POLA1* promoter and a nearby enhancer highlighted predictive subsequences resembling known motifs such as NRF1, SP1, ETS, ATF1, NFY and Inr, some of which exhibited differential contributions to predicted coverage (initiation rates) and profile shape (positioning) (**Fig. 1B**).

We used TF-MODISCO ^90^ to cluster subsequences with high contribution scores across all PRO-cap peaks into a lexicon of non-redundant motif models, each with two complementary representations - a position-weight matrix (PWM) summarizing positional base frequencies and a contribution-weight matrix (CWM) summarizing average contribution scores of each base at each position in the motif, over all its constituent predictive subsequences (**Fig. 2A, columns 1-2**). Discovered motifs matched many well-characterized promoter sequence features, such as two forms of the TATA box and the Inr element (CANT and TANT), the TCT sequence, and other TF motifs associated with promoters and/or active enhancers. Ranking motif CWMs based on their average contribution score to profile shape revealed classical core promoter motifs (the TATA box and Inrs) having the strongest influence on TSS positioning, followed by NFY and the TCT sequence, and then the remaining TF motifs (**Fig. 2A, column 3**).

**Figure 2:**
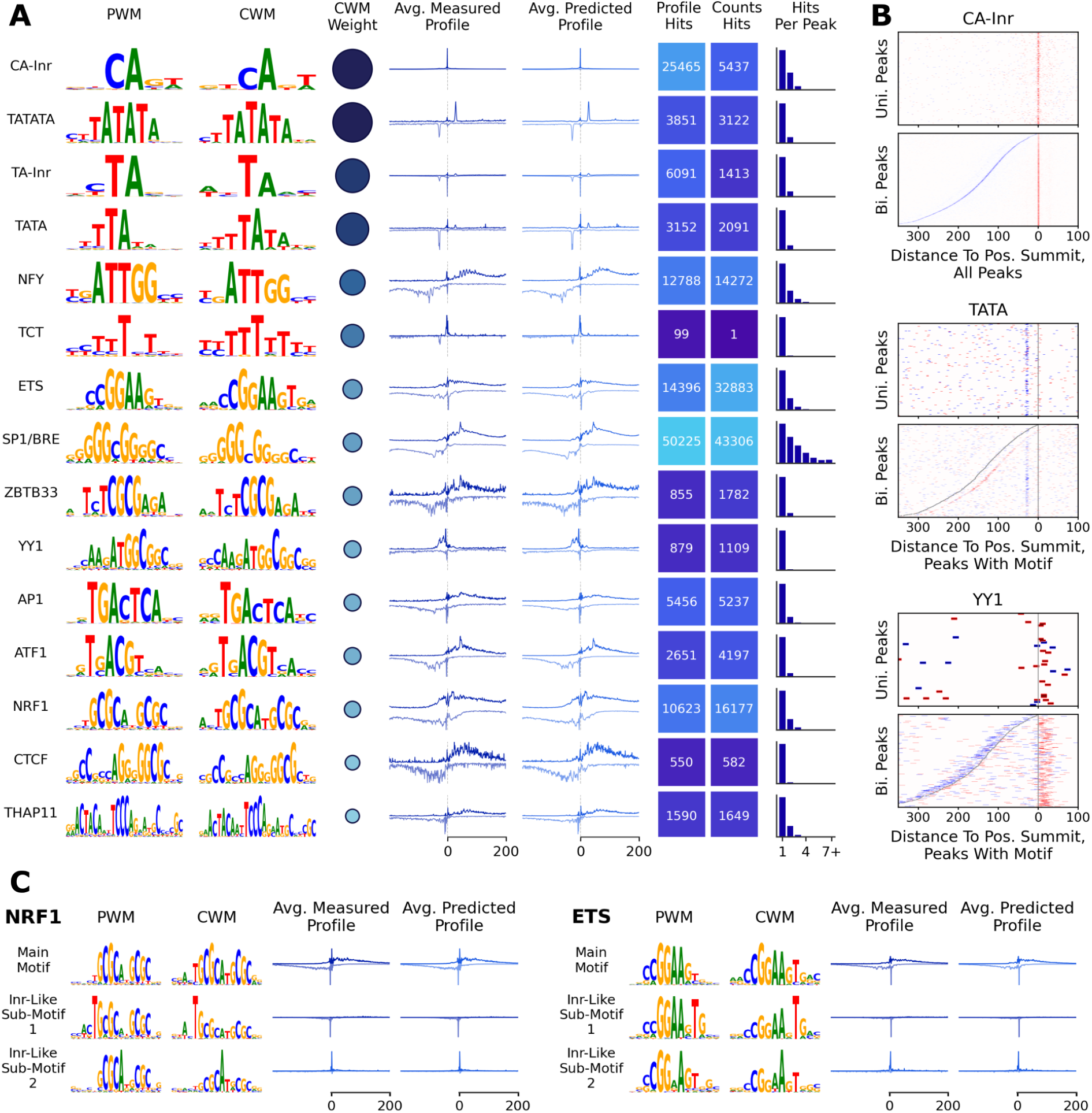
ProCapNet contribution scores highlight a refined set of canonical promoter motifs. (A) Recurrent high-scoring sequence features predictive of initiation identified by TF-MODISCO. PWM: Position-Weight Matrix; CWM: Contribution-Weight Matrix (base frequencies weighted by contribution scores). Columns 1-2 show motifs at normalized heights for visual clarity, while CWM weight (column 3) indicates the magnitude of the motif’s overall contribution, equal to the actual y-axis scale of the CWM. PWMs, CWMs, and CWM weights shown here are from the profile task’s TF-MODISCO output. Average profiles are centered on motif instances, aligned to the same orientation as the PWM/CWM. (B) Identified motif instances relative to PRO-cap peak summits. Uni., unidirectional; Bi., bidirectional (divergent transcription); Pos., positive-strand. Red vs. blue indicates motif orientation, and gray lines indicate PRO-cap summits (omitted from CA-Inr plot because it overlaps the motif hits). (C) Subpatterns found by TF-MODISCO for the NRF1 and ETS motifs that suggest Inr-like secondary roles.

While most TF-MODISCO motifs could be mapped to known initiation elements and associated TF motifs, some esoteric motifs were also discovered. Nineteen esoteric motifs had unusually long (≥15-bp) PFMs with high base specificity across the entire motif, and at least 60% of the sequences supporting each of these these patterns overlapped annotated transposable repeat elements (**Table S1**), which have been previously associated with regulatory activity and TF binding (**Fig. S2A**) ^57,91^. The CWMs of these motifs highlighted shorter segments with high contribution scores suggestive of initiation sequence features embedded within these repeats. A second set of predictive GC-rich repetitive motifs were found downstream of initiation summits, corresponding to where the downstream promoter region (DPR) has previously been identified ^92^ (**Fig. S2B**). Some of these resembled CGG repeats, which have been characterized as structural features in the 5’-UTRs of some genes ^93–96^. No single motif emerged as a definitive marker for this region’s role in initiation, suggesting that the transcription machinery’s preference for the DPR might be non-specific or challenging to represent as a single motif.

We annotated predictive instances of all discovered TF-MODISCO motifs in all PRO-cap peaks as subsequences with high sequence match scores to the discovered motifs as well as high contribution scores ^57^. For all but one motif, over half of the sequence matches were filtered due to low contribution scores, and for 9 of 15 motifs, at least 75% of sequence matches were filtered (**Table S2**). Thus, contribution scores are crucial to identify motif instances that are explicitly used by the models to predict initiation rates and/or positioning, thereby potentially eliminating spurious false positives – especially for short motifs like the Inr ^57^. De-spite the improved filtering induced by contribution scores, we identified predictive motif instances in a vast majority (99.7%, 30,433/30,534) of all PRO-cap peaks and almost all (99.96%, 16,953/16,960) PRO-cap peaks overlapping promoters, substantially improving the coverage of motif annotation of promoters over previous efforts ^22^ (**Fig. 2A**, **columns 6-7**). 28% of the small number (101) of PRO-cap peaks without motif annotations had all reads mapping to only one strand, suggesting they may be artefactual internal recapping events (**Fig. S1E**). PRO-cap peaks on average contained six predictive motif instances that matched on average three unique motifs (**Fig. S2C**). While 96% of peaks contained ten or fewer predictive motif instances, inclusive of homotypic motif multiplicity, 99% harbored instances matching less than seven unique motifs. The most prevalent motifs across all PRO-cap peaks were the SP1/BRE, ETS, CA-Inr, NFY, and NRF1 motifs. These motifs along with THAP11 were also the most likely to have higher motif density in peaks (**Fig. 2A**, **last column**). The rarest motif by far was the TCT sequence, which was identified predominantly in the promoters of ribosomal protein (RP) genes as previously reported ^22^, but was also found at promoters of non-RP genes such as *EEF1A1* and *H3C14-15* as well as at enhancers (**Fig. 1B**).

Next, to understand how initiation events (TSSs) position relative to predictive instances of each motif in PRO-cap peaks, we separately averaged strand-specific measured and predicted PRO-cap profiles centered at all predictive instances of each motif, accounting for motif orientation (**Fig. 2A**, **columns 4-5**). Average observed and predicted profiles were remarkably similar at base resolution, confirming the high predictive performance of the model. The average profiles at both Inr motifs and the TCT sequence indicate that nearly all initiation nearby an Inr/TCT instance occurs directly at the Inr/TCT sequence. Consistent with previous observations, the TATA box shows a strong spacing constraint of ∼30 bp relative to punctate initiation events at TSSs ^40,41,56^ (due to the reverse-complement symmetry of the TATA motif, the orientation of single instances is difficult to disambiguate). The average profiles at most other motifs show broadly distributed downstream initiation and exhibit reverse-complement symmetry, suggesting that the contribution of these motifs may not be orientation-specific. The exception is the YY1 motif, with the bulk of initiation sites in the average profile positioned upstream and biased very strongly towards one strand. We corroborated these positional patterns by also analyzing the positional distribution of predictive motif instances relative to initiation events (summits of peaks) for both unidirectional (single-stranded or sense transcription only) and bidirectional (divergent sense and antisense transcription) PRO-cap peaks (**Fig. 2B**). As expected, the Inr is found precisely at the summits, the TATA box is enriched just around 30bp upstream of summits, and YY1 is enriched within a window about 30bp wide just downstream of summits; all three motifs are strongly biased towards being in an orientation consistent with what strand the PRO-cap signal summit is on. Finally, to quantify the differential impact of different motifs on TSS positioning and initiation rates, we compared the frequency of predictive instances of each motif in all peaks identified using contribution scores relative to the total predicted coverage versus predicted profile shapes (**Fig. 2A, columns 6-7**). Consistent with the previous results, Inr, TCT, and TATA box motif instances influenced TSS positioning more frequently; conversely, ETS, ZBTB33, ATF1, and NRF1 instances influenced initiation rates more frequently.

### ProCapNet reveals cryptic, context-specific initiator codes intertwined within other TF motifs

TF-MODISCO automatically refines each of the reported motifs into higher-resolution submotifs that exhibit subtle differences in base frequencies or contribution scores. We identified several intriguing submotifs of the ETS, YY1, NRF1, AP1, and ATF1 motifs, typically associated with less precise initiation profiles than Inr elements, that appeared to function as potent initiators (**Fig. 2C, Fig. S3**). These submotifs showed higher contribution scores at a subset of bases within the larger TF motif that matched or partially matched the CA-Inr’s consensus sequence CANT (for example, the subsequence AGTG within the ETS consensus sequence CCGGAAGTG). Interestingly, the measured PRO-cap signal at predictive instances of these submotifs was overwhelmingly concentrated directly over the motif, resembling the highly positioned initiation events observed at canonical Inr motifs. The orientation of the Inr-like sub-motifs also correctly aligned with the strand of the initiation spike. For ETS and YY1, the two reported submotifs featured initiation on opposite strands, owing to opposite orientations of two distinct Inr-like subsequences, while for AP1 and ATF1, initiation was only reported on one strand; NRF1 is reverse-complement-symmetric, but has two distinct submotifs highlighting different Inr-like subsequences. These insights from the measured PRO-cap profiles were recapitulated by ProCapNet’s predicted profiles, showcasing the model’s ability to accurately discern the context-specific usage of these cryptic, non-canonical initiation codes intertwined within TF motifs. Thus, ProCapNet reveals that motifs of several TFs can perform dual roles in initiation based on local sequence context, with most canonical motif instances regulating broadly distributed initiation profiles but others acting as cryptic initiators regulating precisely positioned TSSs.

### Promoters and enhancers share a common initiation sequence logic, with differences in initiation driven by differential motif density, diversity, and affinity

Given ProCapNet’s robust prediction performance across both promoters and distal enhancers (**Fig. 1E**), we used the model to investigate whether the two classes of CREs encode shared or distinct initiation sequence logic. First, we compared promoter and distal enhancer sequences based on internal embeddings of the trained ProCapNet model (see Methods). The first two principal components explained 88% of the variance across the embedded space and neatly separated the two classes of elements, indicating that ProCapNet’s latent space is able to distinguish between promoters and enhancers despite not being explicitly trained to do so (**Fig. 3A**).

**Figure 3:**
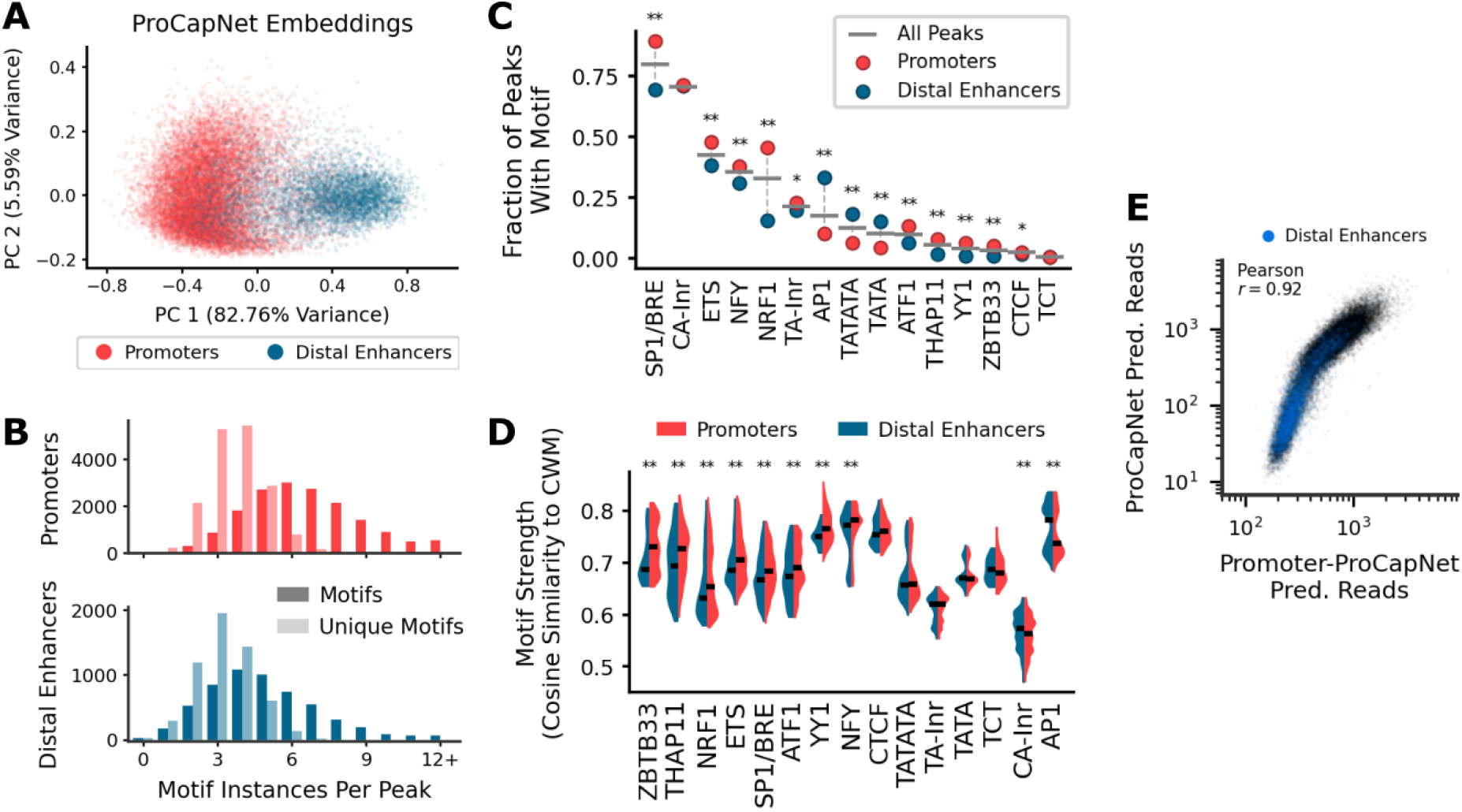
Enhancers and promoters display differential motif complexity. (A) PCA embeddings of internal model representations of sequences at PRO-cap peaks overlapping promoters or distal enhancers. (B) Distributions of the number of identified motif instances in peaks overlapping either promoters or distal enhancers. (C) Fraction of PRO-cap peaks containing at least one instance of a motif, overall vs. in peaks overlapping either promoters or distal enhancers (* = *p <* 0.005, ** = *p <* 10^−19^, two-sided Fisher’s Exact test). (D) Identified motif instance strengths (cosine similarity to the motif CWM) across PRO-cap peaks overlapping promoters and distal enhancers (* = *p <* 10^−2^, ** = *p <* 10^−4^, two-sided Mann-Whitney *U* test). (E) Counts task predictions made on held-out PRO-cap peaks by ProCapNet (y-axis) vs. a re-trained version ProCapNet that only saw promoter sequences during training.

A comparative analysis of predictive motifs instances in PRO-cap peaks in promoters and distal enhancers further revealed that enhancers have fewer initiation-associated motifs and less motif diversity than promoters on average (**Fig. 3B**). Promoters were significantly (*p <* 10^−19^, two-sided Mann-Whitney *U* test) enriched for the SP1/BRE, ETS, NFY, NRF1, ATF1, THAP11, YY1, ZBTB33 motifs and the TANT form of the Inr motif; while enhancers were significantly enriched for AP1 and TATA box motifs (**Fig. 3C**). While similar motif enrichments have been reported using CAGE signal at enhancers ^97^, our analyses suggest additional TF and core promoter motifs. Logistic regression models using motif density or simply motif presence were able to discriminate promoter and enhancer PRO-cap peaks with high (82%) accuracy, further supporting motif density and identity as key properties of initiation syntax differentiating the two classes of elements. Motif instances in promoters were also more likely to have stronger match scores to motif CWMs (which we use as a reasonable surrogate measure of affinity ^98^) compared to those in enhancers, with high affinity instances of ZBTB33, NRF1, ETS, SP1/BRE, and YY1 particularly depleted in enhancers (**Fig. 3D**). Our results indicate that overall lower initiation activity at enhancers relative to promoters can be attributed to a combination of syntactic features including lower motif diversity, density, and affinity, especially of the most common initiator and activator motifs, despite the two classes of elements sharing the same motif lexicon (**Fig. 2A**).

Next, we directly tested whether the *cis*-regulatory logic of initiation learned from promoters would generalize to enhancers. We retrained ProCapNet models only on promoter PRO-cap peaks, using the same cross-validation set up as the original model trained on all types of peaks, and then compared their predictions at all held-out PRO-cap peaks (including enhancers) to those from the corresponding original models (Supplementary Table S3). Profile shape and total coverage predictions from the two models were highly concordant (*r* = 0.88, 0.92 respectively), and approached prediction concordance (*r* = 0.9, 0.98 for shape and coverage) of replicate ProCapNet models trained on identical data with different random seeds. The minor miscalibration of coverage predictions at enhancers between the two models (**Fig. 3E**), as evidenced by the kink in the scatter plot, likely stems from the promoter-only model having to extrapolate PRO-cap signal values outside its training range when predicting signal at enhancers, since the promoters that it was trained on have much higher PRO-cap signals than enhancers on average. These results imply strong generalization of the *cis*-regulatory logic learned by promoter-only models to enhancers.

We then explicitly tested whether the predictive sequence features identified in PRO-cap peaks by the promoter-only ProCapNet models were consistent with those derived from the original models. Similarity (*r* = 0.84) of contribution scores to profile shape predictions from both models across all PRO-cap peaks approached the upper bound (*r* = 0.92) derived from replicate models, suggesting that initiation positioning logic learned at promoters generalizes to enhancers. However, coverage contribution score similarity (*r* = 0.69) was lower, reflecting the miscalibration of total coverage predictions discussed above. Overall, these results suggest that initiation events at promoters and enhancers are regulated by a shared and generalizable *cis*-regulatory code, with higher motif density, diversity, and affinity collectively resulting in stronger initiation at promoters.

### Initiation is regulated by complex *cis*-regulatory logic involving epistatic cooperation and competition across motifs and TSSs

Having established a unified motif lexicon of initiation at promoters and enhancers, we then explored the complexity of *cis*-regulatory logic learned by ProCapNet by asking whether motifs contribute to initiation via independent, additive effects, or whether higher-order syntax-mediated interaction effects play a significant role. An exploratory examination of initiation rates (coverage in *log* scale) relative to total motif counts and unique motif counts showed a non-linear relationship, with coverage saturating at around 10 and 6 motif instances respectively (**Fig. S2D**). A linear model of coverage (*log* scale) as a function of the number of predictive instances of each motif explained a small (*R*^2^ = 0.25) fraction of variance relative to ProCapNet (*R*^2^ = 0.57). Linear models that only used binary presence/absence of predictive motif instances or simply the total number of predictive motif instances in each sequence also fared poorly (*R*^2^ = 0.23, 0.12 respectively). These results suggest that ProCapNet learns more complex regulatory logic than afforded by a simple additive model over motif counts and that the identity of motifs also matters.

Next, using the *MYC* promoter as a case study, we explicitly investigated the interplay between initiation-predictive motif instances at and across multiple TSSs with an *in-silico* motif mutagenesis approach (see Methods) ^57^. The measured and predicted PRO-cap profiles at the *MYC* promoter highlight two distinct TSSs with strong, well-positioned initiation signal on the same strand (**Fig. 4**). Each TSS is ∼30 bp downstream of a predictive TATA box adjacent to an upstream predictive SP1/BRE motif. While the downstream TSS aligns with a predictive canonical TA-Inr instance, the contribution scores under the upstream TSS highlight a putative low affinity Inr instance. ProCapNet also accurately predicts a very weak, cryptic antisense TSS between the two primary sense TSSs, which aligns with a putative low-affinity CA-Inr motif instance containing a C-mismatch at the T in the consensus sequence CANT (reverse-complemented).

**Figure 4:**
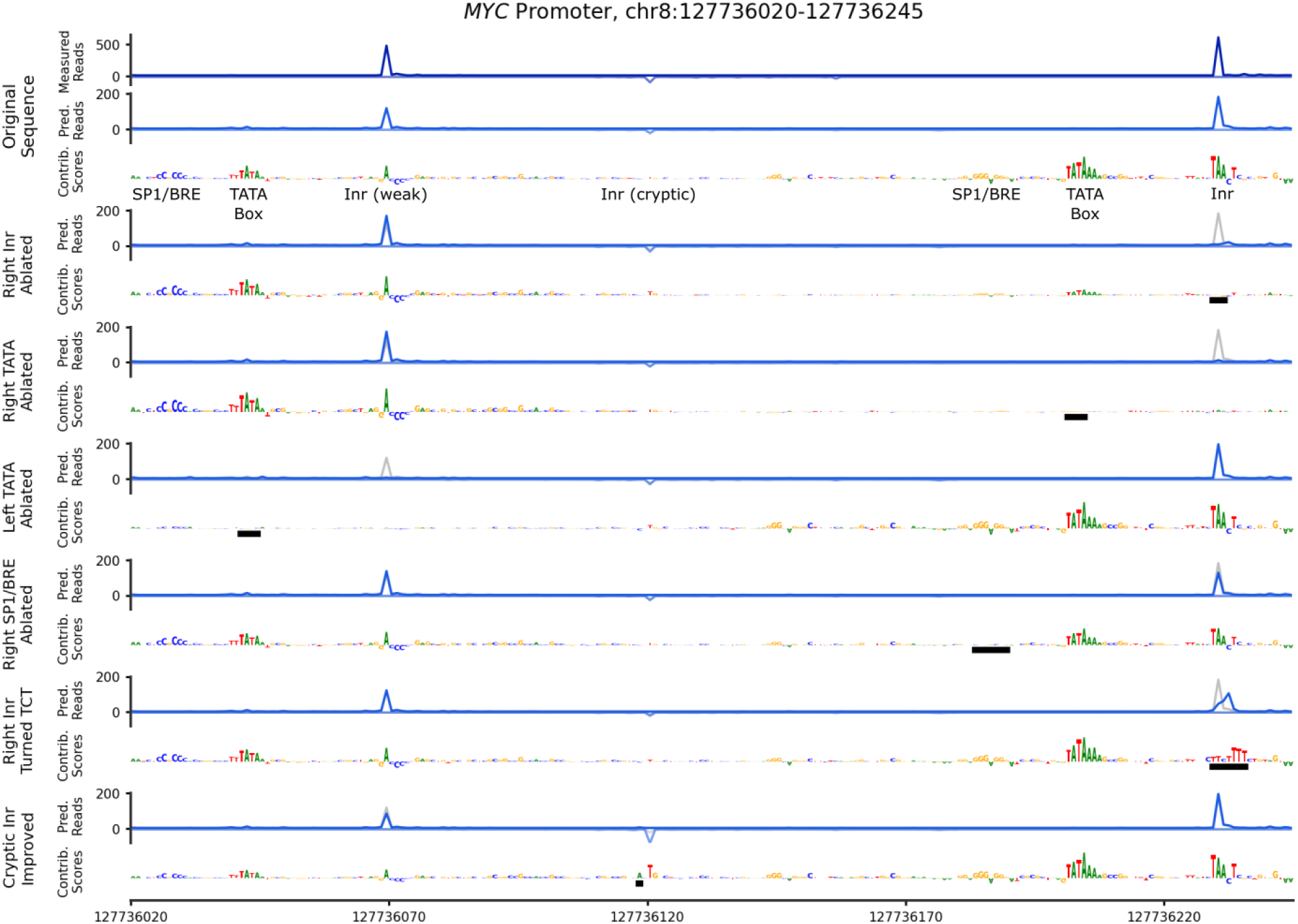
*In-silico* experimentation enables precise investigation of the epistatic interactions between initiation motifs. The measured PRO-cap data and original ProCapNet prediction and contribution scores (profile task) at the primary *MYC* promoter are shown, followed by the model prediction and scores generated from sequences with a single sequence feature modified from the original sequence (indicated by the black boxes). All model predictions, and all score tracks, are plotted on the same y-axis scales.

We used ProCapNet to predict the effects of ablating individual motif instances on nearby initiation signal. We also re-calculated base-resolution contribution scores for each mutated sequence to estimate the impact of motif ablation on contribution scores of other nearby motifs to reveal putative cooperative or competitive motif interactions ^99^ (**Fig. 4**). Ablating the downstream TA-Inr or TATA box resulted in near-complete loss (98% and 95%, respectively) of initiation at the downstream TSS and a dramatic reduction of contribution scores of all other motifs near the downstream TSS, indicative of strong local cooperative interactions between these two motifs that are essential for initiation activity and positioning. Ablating the TATA box near the upstream TSS had an analogous strong localized effect on its activity (97% of predicted initiation lost). In contrast, ablating the SP1/BRE motif near the downstream TSS only partially reduced its activity (by 30%), suggesting a more minor, auxiliary role of the SP1/BRE motif in regulating initiation activity.

Next, we substituted the Inr motif near the downstream TSS with a TCT sequence and observed minor changes to the predicted profile shape at the the TSS without significant loss of predicted local initiation activity (17% decrease within *±*2 bp around the TSS), suggesting some degree of in-context interchangeability between the direct TSS-positioning Inr and TCT motifs.

To examine the potential effects of single nucleotide changes, we corrected a mismatch (G to A) in the low affinity CA-Inr motif at the cryptic antisense TSS, which more than doubled its predicted activity. However, this increased activity at the antisense TSS with the repaired Inr motif was still lower than activity at the two sense TSSs, which have additional TATA and SP1/BRE motifs enhancing local activity.

Having quantified localized effects of motif perturbations, we then investigated whether these perturbations resulted in any epistatic effects across the TSSs in the promoter. Ablating the TA-Inr or TATA box at the down-stream sense TSS resulted in a 44% and 47% increase, respectively, in predicted activity at the upstream TSS. In contrast, ablating the TATA box at the upstream TSS did not impact predicted activity at the downstream TSS. Further, repair of the weak CA-Inr motif at the cryptic antisense TSS which increased its predicted activity resulted in a 30% decrease in predicted activity of the upstream sense TSS, but no change at the downstream TSS. These results suggest a potential complex redistribution phenomenon, where initiation events shift between nearby favorable sites in response to sequence alterations. The redistribution implies asymmetric competition between the TSSs, with the downstream sense TSS being most dominant, likely due to its strong canonical Inr, TATA box, and SP1/BRE motifs.

To comprehensively explore these redistribution effects due to motif and TSS epistasis, we expanded our *in-silico* motif ablation experiments, targeting individual predictive motif instances in all PRO-cap peaks to predict effects on the total coverage and shape of PRO-cap profiles on both strands (**Fig. 5A**; see Methods). Visual inspection of a few candidate loci and motifs highlighted diverse redistribution trends (**Fig. 5B**). For example, Inr ablation often led to loss of initiation at its coincident TSS, while YY1 motif ablation less precisely led to initiation loss within a small window upstream of the motif on the reverse strand, and CTCF motif disruption showed more complex redistribution of initiation from downstream of the motif to up-stream.

**Figure 5:**
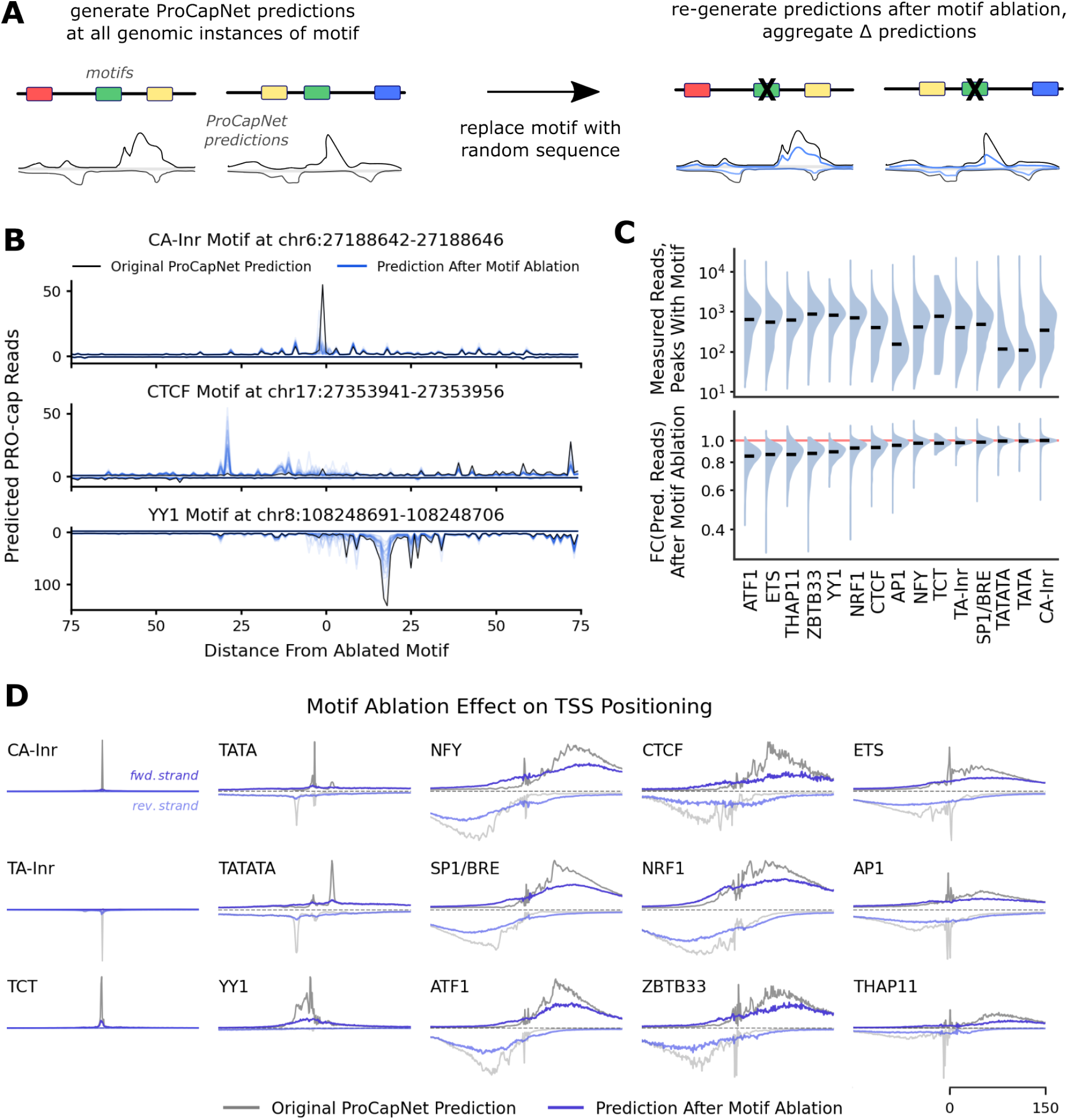
Systematic simulated motif disruption reveals motif contributions to both initiation rate and TSS positioning. (A) Schematic of the motif ablation process. (B) Example loci where individual motif instances were repeatedly ablated and the effect on ProCapNet predictions was estimated for each ablation. (C) The distribution of measured PRO-cap reads across all identified motif instances in PRO-cap peaks (top) vs. the predicted effects of ablating those motifs (bottom). FC, fold-change in predicted reads. Black bars indicate medians. (D) ProCapNet profile task predictions before vs. after motif ablation, aggregated (median) across all instances of each motif. Dashed line indicates where *y* = 0.

We then systematically estimated effect sizes on total initiation activity (fold-change coverage over entire peak region) of all individual instances of each motif in all PRO-cap peaks (**Fig. 5C, bottom**). Inr, TATA box, and TCT motif instances had negligible median effect sizes with relatively tight distributions, reaffirming the primary role of these motifs in localizing initiation at coincident TSSs; the SP1/BRE motif also fell into this category (**Fig. 2A-B**). The resilience of initiation activity to disruption of individual TSS positioning motifs suggests a redistribution of initiation signal from TSSs at the disrupted motifs to other locations within peaks (**Fig. 5B**). In contrast, ablation of ATF1, ETS, and THAP11 motif instances showed the greatest median decrease (∼15%) in total initiation rates, underscoring their roles as consistently strong activators. Notably, for most motifs besides the positioning motifs, effect size distributions were skewed, with long tails towards deleterious effects, particularly for ETS, ZBTB33, and NRF1, indicating that individual instances of these motifs can be quite essential for initiation activity at some PRO-cap peaks. The substantial variation of effect sizes of instances of each motif across peaks further highlights extensive context-dependent epistatic interactions amongst motifs. Juxtaposing the distributions of effect sizes from these counterfactual motif ablation experiments with the observational distributions of measured initiation activity across peaks containing the same motif instances revealed substantial differences in relative ranks of motifs based on their median hypothetical contributions (**Fig. 5C**, **top**). For example, the AP1 motif ranks lower based on the median observational activity compared to counterfactual ablation effects. However, the observational initiation rates in peaks containing AP1 motifs are confounded with the 3-fold higher prevalence of AP1 in distal enhancers, which generally exhibit lower initiation activity than promoters due to more complex motif-driven differences beyond AP1 presence alone (**Fig. 3**). Hence, model-based counterfactual analyses are critical for dissecting the role of motifs while accounting for context-dependent effects on initiation rates.

Next, we assessed the impact of ablating individual motif instances on TSS positioning across entire peak regions. For each motif, we compared median predicted profile probability distributions (normalized by coverage) centered at predictive motif instances before and after their ablation in all peaks, accounting for motif orientation (**Fig. 5D**). As expected, ablating positioning motifs (Inrs, TATA, and TCT) led to precise loss of TSSs at their respective sites; similarly, ablation of the YY1 motif resulted in strand-specific loss of initiation within a 50-bp window upstream. Normalized profile probability predictions (that sum to 1 across 1kb and both strands) imply that a loss in probability density at one location necessitates redistribution elsewhere. However, the absence of concentrated redistributed density in the post-motif-ablation median profiles indicates that these positioning motifs typically don’t shift TSSs to or from any specific location, but rather focus initiation activity at one location that would otherwise be broadly dispersed across the sequence. Other motifs showed broader effects on TSS positioning, generally within *±*150 bp of the motif instance, suggesting a local but less precise influence on initiation sites. The median profiles for these motifs showed reverse-complement symmetry, suggesting they enhance TSS positioning downstream, peaking 50-100 bp away, on both strands evenly. After ablation, these median profiles became flatter, hinting at a general role for these motifs in focusing TSSs with less positional specificity compared to Inrs or TATA boxes. For some motifs, the profile effects indicate a specific redirection of initiation from upstream of the motif on each strand. This effect is most pronounced for the NFY motif: ablation typically leads to both reduced TSS density 50-150 bp downstream and increased density 0-50 bp upstream. This aligns with experimental evidence in mice, showing that NFY loss causes upstream transcription shifts due to altered nucleosome positioning at promoter boundaries ^100^. Similar weaker redistribution effects are also observed for CTCF, SP1/BRE, NRF1, ATF1, and ZBTB33, with a modest increase in predicted TSS density within 50 bp upstream of the motif post-ablation. This suggests that additional TFs may also help focus TSS positioning downstream by mechanisms such as nucleosome positioning that inhibiting initiation upstream. Like NFY, CTCF^101^, BANP (which binds to the ZBTB33/Kaiso motif) ^102^, and ATF1 (in yeast) ^103^ have all been implicated in nucleosome repositioning previously. CTCF has also been implicated in TSS redistribution via blocking of antisense transcription ^104^, which aligns with our observed increase in antisense transcription downstream of the motif upon ablation.

Finally, to understand the relationship between effect sizes of motif disruption on initiation rates versus TSS positioning, we compared the JSD between profile predictions pre- and post-ablation to corresponding fold-changes of predicted initiation rates across all instances of each motif (**Fig. S4**). The correlation between effect sizes on initiation activity and profile shapes varied across motifs (absolute Spearman *r* between 0.05 and 0.42), suggesting that for some motifs, the contributions of a motif instance to positioning and activity are loosely coupled, while for other motifs, individual instances may contribute to both independently.

Collectively, these results suggest that initiation rates and TSS positioning in transcribed regulatory elements are orchestrated by complex, non-additive *cis*-regulatory logic involving cooperative and competitive epistasis across constellations of positioning and activating motifs and associated TSSs.

### ProCapNet distills transcription initiation activity from steady-state RAMPAGE / CAGE expression profiles

The analyses presented above focused on ProCapNet models of nascent RNA PRO-cap profiles because we wanted to isolate the local *cis*-regulatory sequence code specific to transcription initiation, bypassing the need to model additional RNA processing events like Pol II pausing, splicing and degradation, which affect steady-state RNA abundance measured by alternative TSS profiling assays such as CAGE and RAMPAGE ^68,71^. However, since initiation does contribute to CAGE/RAMPAGE measurements, as evidenced by the moderate positive correlation (*r* = 0.67) between PRO-cap and RAMPAGE coverage across PRO-cap peaks in K562 (**Fig. 6A**), we decided to explore what ProCapNet might learn if trained on RAMPAGE profiles.

**Figure 6:**
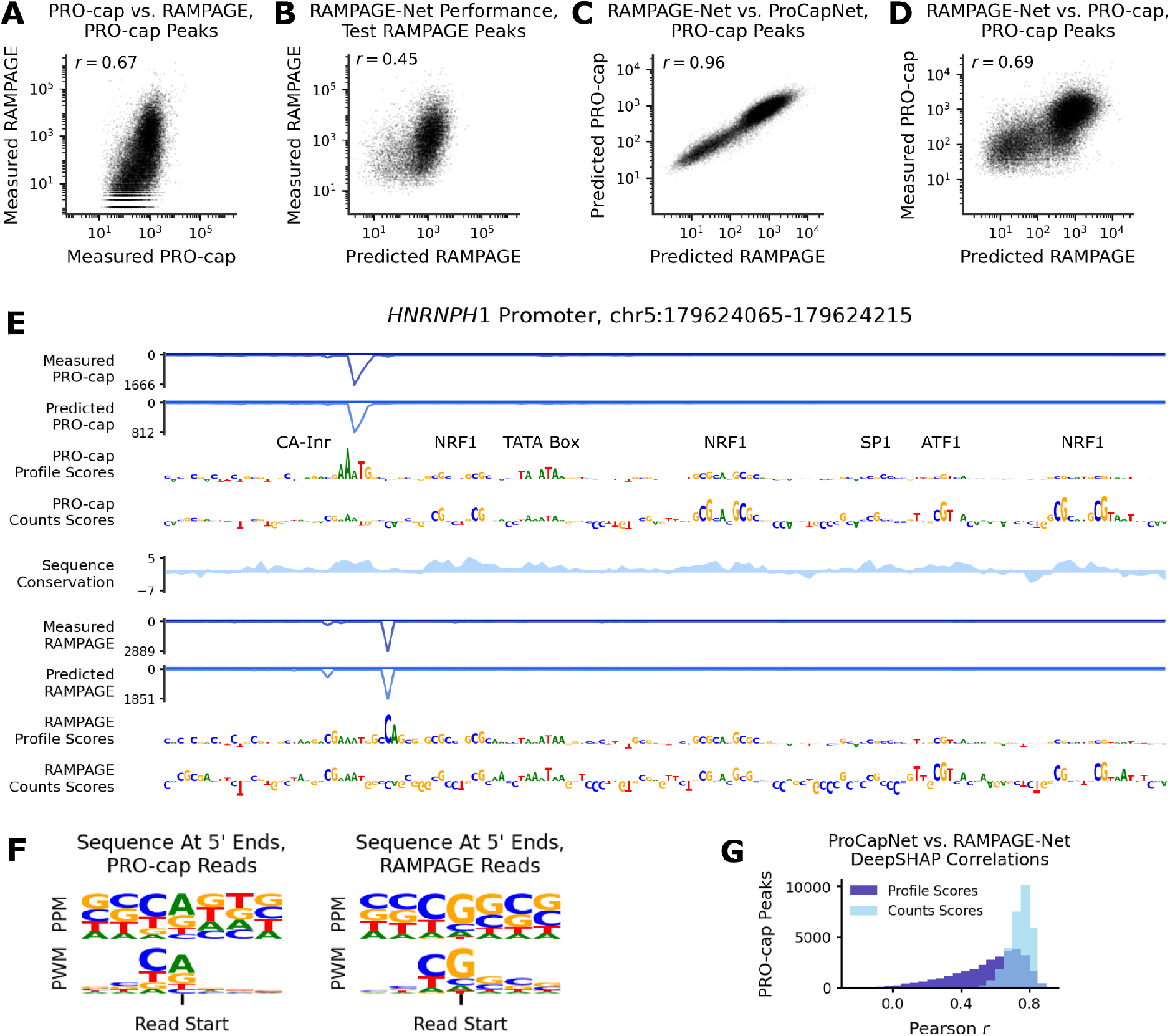
Models trained on RAMPAGE data can learn PRO-cap signal similar to ProCapNet. (A) Relationship between total PRO-cap signal and total RAMPAGE signal within PRO-cap peaks. *r*, Pearson correlation. (B) Counts task performance of RAMPAGE-Net on RAMPAGE peaks from held-out test chromosomes (C) Counts predictions from RAMPAGE-Net vs. predictions from ProCapNet within PRO-cap peaks. (D) Same as C, but with measured PRO-cap signal on the y-axis. (E) PRO-cap and RAMPAGE signal, ProCapNet and RAMPAGE-Net model predictions, contribution scores from both models, and 100-way PhyloP sequence conservation scores ^105^ for an example promoter. (F) Position-probability matrix (PPM) and position-weight matrix (PWM) summarizing all sequences where TSSs (5’ read ends) were identified by PRO-cap (left) or RAMPAGE (right), weighted by the number of reads at each TSS. (G) Pearson correlations between contribution scores from ProCapNet and RAMPAGE-Net for the profile and counts tasks across 1kb windows over all PRO-cap peaks.

Hence, we trained neural networks (called RAMPAGE-Net) directly on base-resolution RAMPAGE profiles from RAMPAGE peaks and accessible background regions in K562, with the same architecture and cross-validation set up as for ProCapNet. RAMPAGE-Net’s prediction performance of total RAMPAGE coverage (*r* = 0.45) and profile shape (normalized JSD = 0.70) at held-out RAMPAGE peaks was notably worse than ProCapNet’s prediction performances at held-out PRO-cap peaks (*r* = 0.71, normalized JSD = 0.45 for coverage and shape respectively), suggesting that steady-state transcription profiles are more difficult to predict than initiation activity from local sequence context (**Fig. 6B**).

Direct comparison of PRO-cap predictions from ProCapNet to RAMPAGE predictions from RAMPAGE-Net at all PRO-cap peaks revealed a remarkably high correlation (*r* = 0.96) between total predicted PRO-cap and RAMPAGE coverage from the two models (**Fig. 6C**), far surpassing the correlation (*r* = 0.67) between measured PRO-cap and RAMPAGE signals. Despite numerous PRO-cap peaks lacking measured RAMPAGE signal (**Fig. 6A**), RAMPAGE-Net imputed RAMPAGE coverage at these regions, with RAMPAGE-Net coverage predictions across all PRO-cap peaks correlating highly with measured PRO-cap (*r* = 0.69) (**Fig. 6D**). These results suggest that the proportion of RAMPAGE signal that RAMPAGE-Net is able to explain from local sequence context is in fact the initiation component of steady-state RNA abundance. Other components of steady-state RAMPAGE signal that relate to post-initiation RNA processing steps, such as splicing and transcript stability, are regulated by sequences further away from the TSS (e.g. splice sites and 3’-UTRs). Hence, the restriction of sequence context local to the TSS appears to implicitly enable RAMPAGE-Net to distill initiation activity *de novo*, on par with ProCapNet trained explicitly on nascent RNA PRO-cap profiles.

However, despite moderate (or high) concordance of measured (or predicted) total coverage between PRO-cap and RAMPAGE, measured (and predicted) PRO-cap and RAMPAGE profile shapes often disagree on precise TSS positioning, as seen at the *HNRNPH1* promoter (**Fig. 6E**). We turned to DeepSHAP sequence contribution scores from ProCapNet and RAMPAGE-Net to their respective profile shape and coverage predictions to resolve the sequence basis of these disagreements. At the *HNRNPH1* promoter, ProCapNet’s profile-shape contribution scores highlight a canonical reverse-complement CA-Inr (ANTG) motif directionally aligned with a strong sense-strand PRO-cap initiation event (putative TSS), which is supported by high PhyloP sequence conservation scores ^105^. In contrast, the RAMPAGE profile suggests a different putative sense TSS, offset from the PRO-cap TSS, aligned with a directionally discordant high scoring CAG motif on the + strand which is not supported by sequence conservation. Thus, the sequence features derived from the models provide more coherent support for the PRO-cap TSS at this locus. Position-frequency matrices of sequences aligned across all TSSs identified by PRO-cap and RAMPAGE respectively also show PRO-cap TSSs most commonly aligning with the Inr sequences CANT and TANT (**Fig. 6F**), matching previously reported Inr consensus sequences ^106^, including adenine as the canonical start nucleotide ^107^, whereas RAMPAGE TSSs favor a non-canonical guanine start. This discrepancy is further reflected in the lower similarity of profile shape contribution scores from ProCapNet and RAMPAGE-Net compared to the similarity of total coverage contribution scores (**Fig. 6G**), which provides a sequence basis for the models’ higher similarity in predicted overall initiation signals over precise TSS positioning. These TSS positioning discrepancies between PRO-cap and RAMPAGE may stem from differences in the experimental protocols. In PRO-cap, the 5’ cap of transcripts is removed prior to adapter ligation, while RAMPAGE retains the G-cap and incorporates additional Gs at the 5’-end during a later template switching step. Sequence features derived from the models provide more convincing support for PRO-cap TSSs than RAMPAGE TSSs.

### ProCapNet predicts similar initiation activity across diverse cell-contexts by learning a shared initiation sequence code with few context-specific features

Our analyses presented above were restricted to ProCapNet models in the K562 cell-line. To understand the generalizability of the models to other cell contexts, we expanded our study by training separate ProCapNet models on additional ENCODE PRO-cap datasets from the A673, Caco-2, CALU3, MCF10A, and HUVEC cell-lines. ProCapNet models from these other cell-lines performed on par with the K562 model for both coverage and profile shape prediction across peaks in each cell-line (**Fig. 7A**). Slight differences in performance metrics across cell-lines likely reflect variations in data quality, exemplified by the highest and lowest coverage prediction correlations being in MCF10A and HUVEC, which also had the largest and smallest numbers of peaks, respectively.

**Figure 7:**
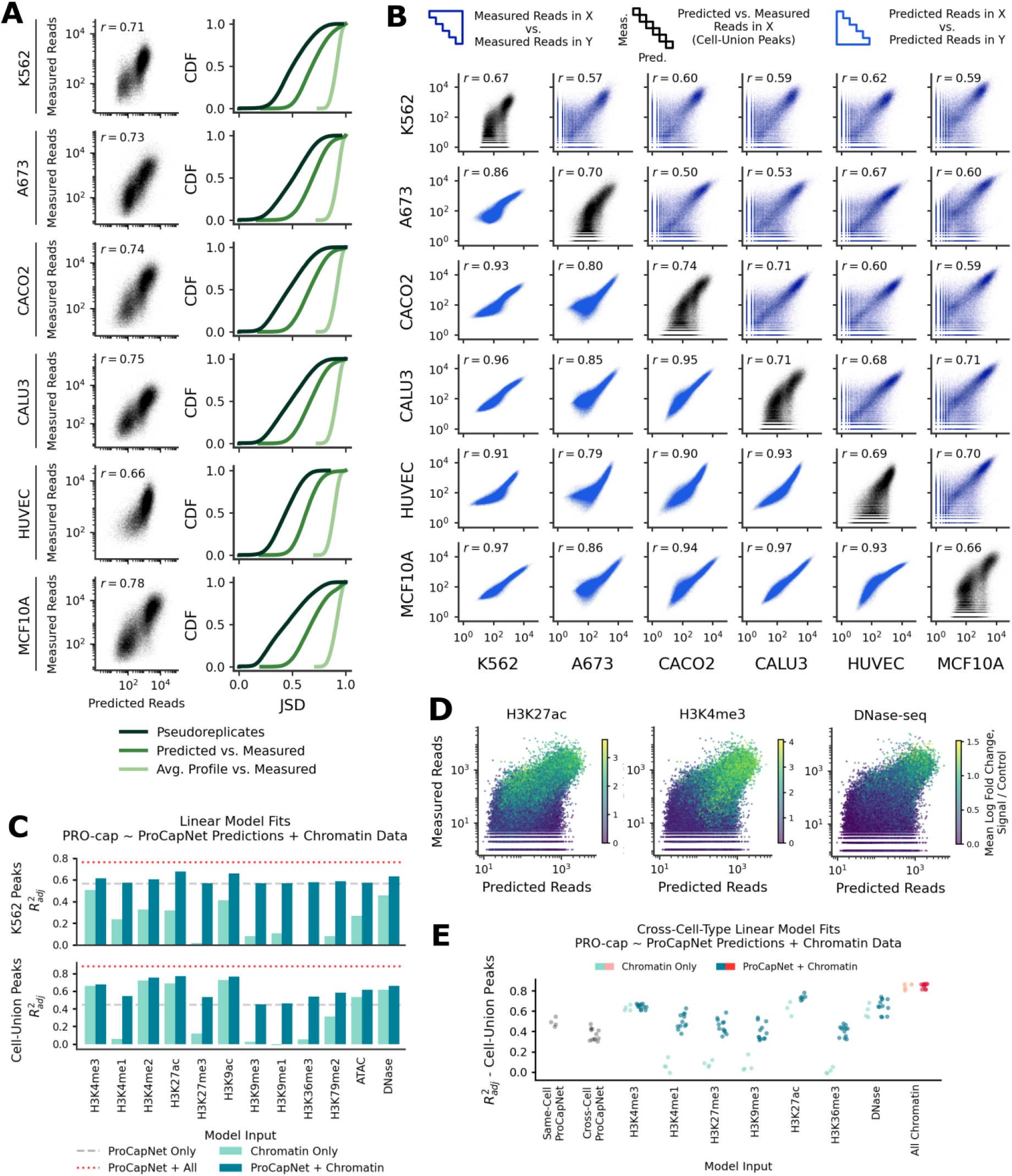
ProCapNet performance and predictions are highly consistent across cell types. (A) ProCapNet performance when trained and tested in all available cell types. (B) Comparison of measured PRO-cap signal (top right triangle) and ProCapNet counts predictions (bottom left triangle) across cell type pairs, plus same-cell ProCapNet predictions vs. measured PRO-cap (diagonal) with peaks active in other cell types included. (C) Goodness-of-fit of linear models of K562 PRO-cap data, using combinations of ProCapNet predictions and various experimental measurements of chromatin state. (D) Examples of how histone modification or DNase-seq signal is distributed across cell-union PRO-cap peaks, with respect to K562 PRO-cap and ProCapNet predictions. (E) Goodness-of-fit of linear models of PRO-cap data in one cell type, using combinations of ProCapNet predictions from another cell type and experimentally measured chromatin state signals.

While these performance evaluations compare measured and predicted PRO-cap profiles across PRO-cap peaks in each cell-line, they do not evaluate the predictions of the model from one cell-line on PRO-cap peaks in other cell-lines. Hence, we first investigated ProCapNet’s ability to predict cell-type-specific TSS positioning in PRO-cap peaks common to multiple cell-lines. We re-examined the *MYC* promoter as a case study, since the downstream TSS showed consistent activity across all cell-lines, while the upstream TSS was uniquely active in K562 (**Fig. S 5A**). ProCapNet models from all cell-lines except K562 accurately predicted the dominance of the downstream TSS as well as the minimal activity of the upstream TSS and the anti-sense cryptic TSS. Profile contribution scores from these non-K562 models also highlighted only the motifs that position the downstream TSS, with no cell-type-specific motifs noted, suggesting potential differential *trans*-regulation of the K562-specific upstream TSS learned by the models. To generalize this observation, for all PRO-cap peaks with at least 50 supporting reads in two or more cell-lines, we tested whether measured PRO-cap profile shapes in each cell-line were more similar to predicted profile shapes from ProCapNet models of the same cell-line compared to predictions from other cell-lines. Measured TSS profiles correlated better with predicted TSS profiles from the same cell-line than predictions from other cell-lines 80% of the time, confirming ProCapNet’s ability to learn cell-type-specific TSS positioning. These results also emphasize the importance of training cell-type-specific models to discern cell-type specificity of TSSs, and highlights a significant limitation of previous cell-type-agnostic models trained on profiles averaged over diverse cell types ^77^ (**Supplemental Note**).

Next, we assessed if ProCapNet models from the diverse cell-lines could also accurately predict cell-type-specific initiation activity. For each pair of cell-lines, we compared measured coverage and predicted coverage across the union of peaks from all cell-lines. For all cell-line pairs, the correlation between predicted PRO-cap coverage (*r* = 0.8−0.97) (**Fig. 7B**, **lower triangle**) was much higher than the correlation between measured coverage (*r* = 0.5 − 0.71) (**Fig. 7B**, **upper triangle**), suggesting that ProCapNet models trained in different cell-lines make largely cell-type-invariant predictions of initiation activity. ProCapNet models trained in each cell-line showed a predisposition to predict initiation activity at regions that are inactive in that cell-line but active in others (**Fig. 7B**, **diagonal**), despite accurate predictions of activity across peaks within each cell-line (**Fig. 7A**).

To further investigate the sequence determinants of these largely cell-type-invariant initiation activity predictions from models trained in different cell-lines, we compared TF-MODISCO motifs derived from each cell-line’s models. All K562 motifs were identified in all other cell types, except for the TCT sequence, which was found in 4 of the 6 cell-lines, and the TATA box, for which the two subtypes were merged into one in half of the cell-lines (**Fig. S5B**). Motif prevalence was also highly consistent across all cell-lines. Some cell-line-specific motifs were also identified, including some ubiquitous nuclear factors and some cell-type-specific TFs. For example, the motif for EWS-FLI, a marker oncogenic factor specific to Ewing sarcoma, was exclusively found by TF-MODISCO in the A673 Sarcoma line ^108^ (**Fig. S5C**). Similarly, motifs of HNF4 and FOX TFs, which have specific functional roles in colorectal tissue, were exclusively found by TF-MODISCO in the Caco-2 colorectal cancer cell-line ^109–111^. Hence, ProCapNet models trained in different cell-lines learn a predominantly shared *cis*-regulatory code and a few cell-type-specific motifs, resulting in largely cell-type-invariant predictions of initiation activity.

### Cell-type specificity of initiation activity is strongly influenced by local chromatin state

ProCapNet’s predictive motifs often exclude key regulators of cell-type-specific chromatin accessibility and active histone modifications, such as the motifs of GATA1 and TAL1 in K562^112,113^. This decoupling of sequence determinants of initiation from chromatin state might stem from our training set design, which included PRO-cap peaks and accessible, but transcriptionally inactive background regions from each cell-line. But it is more likely to be a consequence of ProCapNet’s exclusion of distal regulatory sequences, crucial for accurate prediction of cell-type-specific chromatin state, and, by extension, cell-type-specific initiation activity ^62,63^. To further understand the relationship between chromatin state and initiation, we tested whether explicit incorporation of various chromatin state markers might improve prediction of initiation activity and its cell-type specificity.

We examined how ProCapNet predictions relate to chromatin state by fitting linear models to initiation activity in K562 using combinations of ProCapNet’s initiation activity predictions, measured chromatin accessibility, and/or ChIP-seq histone modifications in K562 as inputs. We report the adjusted 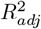 of the linear models, which corrects for the number of input predictors and training data points. We first tested this strategy using PRO-cap peaks in K562 and later expanded to a combined set of peaks from all cell types.

Restricting to PRO-cap peaks in K562, linear models using ProCapNet predictions alone had an 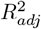 of 0.57, while a linear model using all chromatin markers achieved an 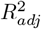 of 0.71. Combining chromatin markers with ProCap-Net predictions elevated the 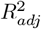 to 0.73 (**Fig. 7C, top**). Thus, chromatin state is strongly associated with initiation activity in active PRO-cap peaks. The chromatin markers with the strongest marginal association with PRO-cap activity were H3K4me3, H3K9ac, and DNase-seq, consistent with prior studies (**Fig. 7C, top, light bars**) ^48^. However, no individual chromatin marker outperformed ProCapNet alone.

Next, to assess which chromatin markers complement the local initiation logic learned by ProCapNet, we fit linear models to predict measured PRO-cap using both individual chromatin markers and ProCapNet predictions as input. Chromatin markers with stronger marginal association with PRO-cap activity did not always correspond to greater improvement over ProCapNet alone (**Fig. 7C, top, dark bars**). H3K27ac and H3K9ac were the most complementary, followed by DNase-seq, H3K4me3, and H3K4me2. Other histone modifications and ATAC-seq showed minimal additional improvement over ProCapNet predictions alone. Further, no single chromatin marker, when combined with ProCapNet, matches the performance of all of them combined.

Expanding the analysis to PRO-cap peaks active in any of the cell-lines clarified the importance of local chromatin state for cell-type-specific initiation activity in K562. The most complementary chromatin signals in K562, identified in the above analysis, each showed distinct patterns of higher signal in PRO-cap peaks active in K562 compared to peaks inactive in K562 but active in other cell-lines (**Fig. 7D**). Linear models trained to fit PRO-cap activity in K562 from different combinations of inputs over this expanded set of PRO-cap peaks showed that models based on ProCapNet predictions alone had substantially lower 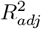 (0.45) than models that included all chromatin markers (0.86; **Fig. 7C**, **bottom**). For all chromatin markers except H3K4me1, H3K9me3, and H3K9me1, marginal associations with measured PRO-cap reads were also greater than associations estimated over K562 PRO-cap peaks. Further, most linear models trained on the expanded peak set that combined ProCapNet predictions with individual chromatin markers had higher 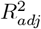 than a model using ProCapNet predictions alone (**Fig. 7C**, **bottom, dark bars**). H3K9ac and H3K27ac were top contributors, followed by H3K4me2, H3K4me3, and chromatin accessibility; these models only marginally outperformed models that only used chromatin markers (**Fig. 7C**, **bottom, light bars**).

Finally, for each pair of cell-lines, we tested how well integrating chromatin markers in a target cell-line with ProCapNet predictions from models trained on a different reference cell-line could fit PRO-cap activity in the target cell-line by adjusting model predictions for cell-type-specific peaks. We performed this analysis for all cell type pairs for which shared chromatin marker datasets were available (K562, A673, Caco-2, and HUVEC). First, ProCapNet predictions were less accurate in fitting measured PRO-cap signals when the model was trained in a different cell type, confirming that ProCapNet captures some degree of cell-type specificity (**Fig. 7E**). While all target-cell-line chromatin-augmented ProCapNet models showed improvements over using ProCapNet alone, the extent of improvement varied significantly depending on the chromatin markers incorporated and the cell-line pairs. H3K4me3, H3K27ac, and DNase-seq provided the largest gains in 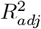, whereas H3K27me3, H3K9me3, and H3K36me3 provided marginal improvements. Models that combined reference ProCapNet predictions with all chromatin markers from the target cell-line had the best fits (average 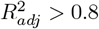), generally outper-forming models that used any single marker, although models augmented with H3K27ac alone were marginally worse than the model augmented with all markers.

Collectively, these results suggest that local chromatin state is critical for predicting cell-type-specific and differential initiation activity at regulatory elements across cell types and less relevant for predicting initiation across regions that are active in a specific cell type. ProCapNet learns a *cis*-regulatory code of initiation that partially complements cell-type-specific sequence drivers of local chromatin state. Hence, ProCapNet’s predictions tend to be more consistent across cell types without explicit integration of cell-type-specific chromatin state.

## Discussion

ProCapNet is a compact neural network based on the versatile BPNet architecture that effectively models transcription initiation rates and positioning of initiation events from sparse, base-resolution PRO-cap profiles, using only local DNA sequence context. Despite the significant influence of distal regulatory elements on transcription, ProCapNet’s effectiveness at predicting initiation profiles suggests that local sequence context encodes a substantial component of the *cis*-regulatory logic of initiation.

Querying ProCapNet models with a suite of well-established, robust model interpretation methods enabled identification and systematic analysis of *cis*-regulatory sequence features and their higher-order syntax that predict initiation across the human genome, addressing several crucial questions about the role of sequence in transcription initiation regulation in *cis*.

ProCapNet’s multi-scale approach to modeling initiation rate and profile shapes separately enables delineation of the specific contributions of both known and novel variants of initiation motifs and general transcription factor (TF) motifs to initiation rate and transcription start site (TSS) positioning, both individually and in combination. While some motifs primarily specify TSS positioning, others modulate initiation activity more broadly, and some influence both aspects depending on the sequence context. Each motif contributes uniquely to TSS positioning and can shift initiation upstream or downstream of its location.

These findings align with several previous observations of the role of different TFs and motifs in initiation. First, earlier work has proposed that bidirectional, divergent transcription is the product of two opposite-oriented core promoter motif sets ^80,114,^ ; indeed, ProCapNet attributes its accurate predictions in divergent PRO-cap peaks to opposite core promoter motif sets, providing supervised confirmation of this model (**Fig. 2B**). Second, Core *et al*. 2014^48^ described two groups of TFs that bind within divergent TSS pairs: central-binding factors, responsible for overall activation of transcriptionally active regions, and TSS-proximal TFs, which include GTFs and which bind much closer to where initiation events are positioned; interpretation of ProCapNet revealed motif contributions that can be cleanly categorized into these two groups, with the latter describing core promoter motifs and YY1, and the former describing the remaining TF motifs. Third, TATA boxes, Inrs, and TCT sequences have been implicated in TSS positioning previously ^40,56^. ProCapNet precisely quantifies the positioning contributions of these motifs, but beyond that, it represents a unified model of how TATA boxes, initiator elements, and other TF motifs combinatorially contribute to positioning, including at regions of dispersed transcription, where TATA boxes and other motifs are less common ^22^.

While ProCapNet robustly recovered motifs traditionally associated with transcription, such as the TATA box and Inr element, it provided limited evidence for the GC-rich DPR element as previously reported ^92^. Although ProCapNet highlighted general GC-rich features downstream of TSSs, that no single representative motif was discovered suggests that the contribution of the DPR may be more general than a lone PWM could effectively represent (i.e., the sequence composition of the DPR does impact initiation, but there is no discrete DPR motif). The DPR could also play more of a role in downstream regulatory processes, such as promoter-proximal pausing, RNA stability, splicing, etc. than in initiation. In particular, because the region downstream of the TSS is by definition transcribed, it might influence abundance measurements by RNA-seq or CAGE/RAMPAGE more than nascent transcription read-outs such as PRO-cap. Hence, predictive sequence models such as ProCapNet trained directly on nascent RNA initiation profiles may be better suited to characterizing the role of the DPR and any other motif in initiation specifically.

ProCapNet also highlighted the surprising ability of specific instances of the NRF1, ETS, YY1, AP1, and ATF1 motifs to act as context-dependent initiators via cryptic Inr sequences intertwined within these motifs. This dual functionality has been previously reported for YY1 and ETS, but not for NRF1, AP1, or ATF1^38,116^. The alternative initiator functionality of these motifs is predictable from sequence alone, yet the sequence of the TF motif instances that directly initiate transcription are not conspicuously different from those of their canonical counterparts, suggesting that local sequence context of motif instances is crucial in determining when this secondary functionality is activated. This finding raises several mechanistic questions, including whether or not the TFs that typically bind these motifs also bind to instances that function as Inrs, whether this binding influences the recruitment of general transcription factors, and whether these TFs remain engaged with the promoter when components of the pre-initiation complex, which interact directly with Inr elements, are assembled (as has been shown for YY1^38^). One potential model is that an equilibrium between binding of the TF and binding of the PIC at these sites serves to maintain accessibility of the core promoter. This set of TFs also overlaps with those previously identified as having an initiation-centric role in promoters that are less responsive to enhancers (primarily housekeeping genes), suggesting a potential role for dual-initiator functionality in enhancer-promoter compatibility ^52^.

ProCapNet improved genome-wide annotations of motifs involved in initiation across all classes of transcribed regulatory elements. We detected at least two initiation motifs in nearly every active promoter in the K562 PRO-cap datasets and in 98% of PRO-cap peaks overall. This finding significantly revises previous estimates of initiation relevant motifs in promoters and reinforces the view that local sequence context encodes a substantial component of the *cis*-regulatory logic that regulates initiation at all human promoters. Differences in nascent transcription between promoters and enhancers could be clearly attributed to syntactic variations in motif density, diversity and affinity, rather than distinct motif lexicons, supporting the hypothesis that enhancers and promoters form a functional continuum i.e. many enhancers, in addition to influencing distal promoter activity, also act as weak promoters on their own ^47,48,1^.

Systematic *in-silico* motif ablation studies with ProCapNet models at individual loci and genome-wide highlighted complex, modular rules of epistatic motif syntax and competitive interactions among transcription start sites (TSSs), with significant predicted redistribution of initiation activity upon motif perturbations. These model-based counter-factual analyses proved critical for dissecting the role of motifs, accounting for context-dependent effects on initiation rates and positioning. Overall, these findings emphasize the extensive intricate interplay of cooperative and competitive mechanisms among motifs and TSSs, shaping the transcription initiation landscape across promoters and enhancers. These results especially highlight the advantage of supervised deep learning models that can learn *de novo*, nonlinear predictive representations and contextual epistasis from raw sequence, relative to traditional computational approaches for sequence analysis that rely on unsupervised statistical over-representation of individual features often confounded by extensive non-causal correlations between features.

ProCapNet’s architecture can also be used, as-is, to model alternative TSS profiling assays such as CAGE and RAMPAGE that measure steady-state mRNA abundance. Despite the significant contribution of RNA processing to RAMPAGE signals, which cannot be learned from local promoter sequence context, these RAMPAGE-Net models effectively distill out the initiation portion of the signal driven by promoter syntax, which is remarkably concordant with initiation activity predictions from ProCapNet models trained on PRO-cap data. Hence, initiation activity attributed to local sequence and the underlying sequence syntax could be potentially imputed from widely available RAMPAGE or CAGE data in diverse cell contexts ^119,120^. However, comparative analysis of performance and interpretation of RAMPAGE-Net and ProCapNet models helped us identify systematic differences in TSS precision between RAMPAGE and PRO-cap measurements potentially linked to experimental or mapping artifacts in the RAMPAGE experiments we modeled. Hence, interpretable sequence models also serve as a powerful lens to decipher subtle idiosyncrasies present in different experimental assays.

Predictive sequence models trained on initiation profiles could inadvertently learn sequence drivers of upstream processes, such as TF binding and chromatin accessibility, that indirectly influence initiation. To focus ProCapNet on sequence determinants of initiation decoupled from these upstream processes, we explicitly included during training transcriptionally inactive but accessible background regions from the training cell-type context. Subsequent interpretation of ProCapNet models trained in diverse cell-lines showed that they often do not learn key cell-type-specific drivers of chromatin accessibility (e.g. GATA1 in K562) and instead learn a partially complementary *cis*-regulatory code of initiation that is largely cell-type-invariant. ProCapNet models learn motifs exclusive to initiation (e.g. TATA boxes and Inrs), as well as TF motifs (e.g. NRF1, YY1, and AP1) that also influence accessibility, but in a cell-type-invariant manner ^112^. However, cell-type-specific motifs (e.g. EWS-FLI in A673, HNF4 and FOX in Caco-2) are occasionally learned by ProCapNet, raising interesting questions about the precise mechanism by which these rare cell-type-specific motifs influence initiation. The largely cell-type-invariant code learned by ProCapNet results in imperfect prediction of cell-type-specific initiation activity. Post-hoc integration of ProCapNet with chromatin accessibility and histone modifications, individually or collectively, resulted in substantial improvement of cell-type-specific predictions, highlighting the critical role of chromatin state for cell-type-specific initiation activity ^48,78^. These results emphasize the importance of training set design, complementary performance evaluation strategies, and thorough model interpretation for developing models that balance interpretation goals with prediction performance, and to identify strategies to enhance both. Our findings also suggest that initiation assays and associated predictive models like ProCapNet will likely improve prioritization and interpretation of noncoding variants that disrupt sequence syntax exclusive to initiation. A recent study already demonstrates that ProCapNet complements analogous models of chromatin accessibility ^112^ and outperforms a state-of-the-art, long-context sequence model of gene expression ^62^ at predicting and explaining the effects of engineered sequence edits in enhancers and promoters on gene expression ^121^.

ProCapNet offers several advantages over the recent Puffin model, which also aims to predict base-resolution PRO-cap and CAGE profiles averaged over diverse biosamples from local sequence ^77^ (**Supplemental Note**). Puffin is constrained to learn a restricted set of motifs curated from a larger, long-context model using an ad-hoc, iterative distillation approach. Puffin also uses an explicitly additive neural network architecture, based on an unverified assumption that motif instances impact initiation independently and additively. Puffin emphasizes this hand-crafted, transparent model design for direct interpretation of model parameters, suggesting that it enhances the discovery of initiation syntax. We systematically benchmarked ProCapNet against Puffin to understand their pros and cons (**Supplemental Note**). First, we found that ProCapNet outperforms Puffin at prediction, benefiting significantly from being a cell-type-specific model. Second, all motifs and their positional preferences identified by Puffin are independently discovered by ProCapNet (except the U1 snRNP, which is a post-initiation RNA processing regulator). However, ProCapNet discovers several important properties of the initiation code missed by Puffin due to its constrained design. Puffin does not identify several ProCapNet motifs predicted to influence initiation including ubiquitous motifs such as CTCF and several cell-type-specific motifs (**Fig. S 5B**). Puffin’s additive model does not accommodate motif epistasis, severely limiting its ability to accurately identify active instances of Inr motifs and decipher cryptic, context-specific initiator codes intertwined within other TF motifs. The critical role of motif epistasis in initiation as highlighted by ProCapNet is strongly supported by other recent independent computational and experimental studies ^122,123^. Furthermore, while Puffin’s training scheme is based off the assumption that aggregating data across many cell types is necessary to learn the relationship between sequence and initiation, ProCapNet’s accuracy refutes that claim, and we show how data aggregation can even lead to misleading model predictions. Unlike Puffin, ProCapNet models and interprets sequence drivers of overall initiation activity levels in addition to TSS positioning. Finally, our study provides a unified analysis of the *cis*-regulatory code of initiation at all transcribed regulatory elements, including enhancers, while Puffin is restricted to promoters. In summary, the Puffin model does not offer any advantages over well-established post-hoc interpretation methods applied to unconstrained, black-box architectures such as ProCapNet, and in fact hinders the ability to discover important syntactic properties of the initiation code.

Going forward, ProCapNet can be extended to address some key limitations. First, our current study explicitly focuses on quantifying the influence of local sequence context on initiation activity and positioning and dissecting the influence of predictive local sequence features and their higher-order syntax. While ProCapNet can explain a substantial component of initiation activity, it will be critical to incorporate the influence of distal regulatory elements to further improve predictive performance and cell-type specificity. However, the key challenge is to design long-context models that incorporate these enhancements without sacrificing model stability and interpretability ^62–64^. Second, incorporation of sequence drivers of chromatin state will also be critical to improve model performance. While our current integrative models incorporate experimental measures of chromatin state, future efforts could jointly train and interpret sequence models of chromatin accessibility, histone marks, and initiation activity to enhance prediction performance while preserving modular interpretability of sequence drivers of the different readouts ^112^. Third, while we focused on isolating regulation of initiation alone, initiation is not independent of promoter-proximal pausing in humans ^67^; even PRO-cap measurements of initiation activity can be underestimates of the initiation capacity of a promoter if pausing is the true limiting step. Thus, it’s possible that the model’s overall activity predictions could be improved by taking pause rates at individual loci into consideration. Finally, ProCapNet has enabled robust discovery of several novel hypotheses about the *cis*-regulatory code of initiation at individual loci and globally; experimental validation will be critical to verify them.

Nevertheless, ProCapNet and its future extensions could be used to address several other interesting questions, such as the subtle differences between specific classes of regulatory elements, including promoters of housekeeping vs. cell-type-specific genes and lncRNAs. These models could be used to improve the base-pair precision of transcription start site annotations, systematically screen for non-coding genetic variants in rare and common diseases that impact transcription initiation in diverse cell contexts, and study the evolution of the *cis*-regulatory code of initiation across species. The models could also be used to optimize design of synthetic promoters, benefiting experimental reporter library designs and gene therapy payloads. Finally, sequence models of initiation coupled with analogous models of TF binding, chromatin accessibility, histone modifications, RNA processing, stability, and abundance could reveal the complex interplay between partially overlapping layers of the *cis*-regulatory code of transcriptional regulation. As these models evolve, they will continue to enhance our understanding of the genome, human biology, and disease.

## Methods

### PRO-cap datasets

Uniformly processed PRO-cap peak calls and UMI-filtered read alignments for all replicate experiments were downloaded from the ENCODE portal for all cell-lines: K562 (accession ID ENCSR261KBX) for the main ProCapNet model, as well as A673 (ENCSR046BCI), Caco-2 (ENCSR100LIJ), Calu3 (ENCSR935RNW), HUVEC (ENCSR098LLB), and MCF10A (ENCSR799DGV) for the additional cell-line models ^124^. Unidirectional and bidirectional peak calls were combined into a single peak set. The first read in read pairs was filtered out of the read alignments, and only the single base at the 5’ end of the second read (corresponding to the 5’ end of nascent RNAs, or the TSS) was retained as a data point for each read. Data was then merged across replicates but kept separate by strand.

### Other datasets

All other experimental datasets were downloaded from the ENCODE portal ^124^. For model training, peak calls of DNase hypersensitive sites for each cell-line were obtained from accession IDs ENCSR000EKS, ENCSR789VGQ, ENCSR114QAK, ENCSR255STJ, ENCSR366NBE, ENCSR000EOQ for K562, MCF10A, A673, Caco-2, Calu3, and HUVEC, respectively.

For analysis of model performance and motif instances across candidate *cis*-regulatory elements (cCREs), we downloaded cell-type-specific ENCODE annotations (ENCSR301FDP) ^79^.

For analysis of quantitative chromatin state, we used processed ENCODE data: specifically, fold-change over control ChIP-seq tracks for histone modifications, read-depth normalized signal tracks for DNase-seq, and fold-change over control tracks for ATAC-seq. ENCODE accession IDs for most experiments are in Supplementary Table S4; we also used the following datasets only available in K562: ATAC-seq (ENCSR868FGK), H3K4me2 (ENCSR000AKT), H3K79me2 (ENCSR000APD), H3K9ac (ENCSR000AKV), and H3K9me1 (ENCSR000AKW).

For RAMPAGE data (ENCSR000AER), we obtained base-resolution experimental signal by taking the read 1 5’ ends from the alignment bam files and merged the “transcription start site” peak calls from each replicate to create our peak set.

The GRCh38 genome sequence and gene annotation (v41) were downloaded from GENCODE ^125^. For mappability-aware training and assessment of how model performance varies with mappability, we used the k36 multi-read hg38 Umap track downloaded from https://bismap.hoffmanlab.org/ ^126^. Tracks of 100-way PhyloP sequence conservation ^105^ were obtained from the UCSC genome browser ^127^.

### ProCapNet model design

ProCapNet architecture is adapted from the BPNet model, previously described in Avsec *et al*. 2021^57^. ProCapNet takes as input 2,114 bp of sequence and outputs 1) a vector representing the probability of an initiation event being observed at each base on both strands within a 1,000-bp window and 2) a scalar representing the total *log*-count of initiation events (PRO-cap reads) within the same window (Figure 1A). ProCapNet differs from BPNet in the following ways. First, while BPNet’s predictions are conditioned on an experimental control track, ProCapNet does not integrate information from experimental controls, since none exist for PRO-cap. Second, ProCapNet’s sequence input is of size 2,114 bp and the output is 1 kb, whereas BP-Net’s input sequences and output predictions are both 1kb wide; ProCapNet’s increased input size removes the need to pad input sequences with zeros to allow convolutional filters to scan at the sequence edges. This change avoids any artefacts at the borders of predictions arising as a result of zero-padding.

Third, the ProCapNet counts task predicts the *log* of the number of PRO-cap reads with 5’ ends mapping within a 1,000-bp window, summed over both strands, while BP-Net makes one prediction per strand. A similar modification is made to the profile task: while the profile heads of both models make base pair-resolution predictions across both strands, BPNet treats the strands as two independent prediction tasks and applies a softmax to each strand individually, whereas ProCapNet applies the softmax function across the predictions for both strands, as if they are one array. These modifications to strand representation allow the ProCapNet profile task to learn to predict asymmetry in read allocation across strands, while the counts task predicts overall rate of initiation across either strand. ProCapNet is trained using the same loss functions as BPNet for both profile and counts tasks (MSE and a multinomial loss, respectively), but because the predictions are represented as combined across strands for ProCapNet, each loss is calculated once per input sequence, rather than once per strand per input sequence.

Fourth, ProCapNet implements a mappability-masked training scheme, incorporating information about which bases are not uniquely mappable by PRO-cap-length sequencing reads into the profile task loss function. Specifically, any base that is not entirely uniquely mappable, according to 36-mer multi-read annotation tracks produced by the Umap software ^126^, is assigned a loss weight of zero, to avoid penalizing the model for any incorrect predictions at bases at which the measured PRO-cap data may be inaccurate. This approach is only implemented during training; all reported performance metrics include performance on bases that are not uniquely mappable. Models trained with this approach had extremely similar performance to those trained without it, but contribution scores showed mild qualitative improvement.

Finally, all post-training ProCapNet predictions were generated by taking the average of the model output for both the forward strand sequence and its reverse complement, to increase robustness of predictions ^128^.

The following model architecture hyperparameters were tuned to optimize performance, according to the validation set for the first fold K562 model:

1. Eight dilated convolutional layers (9 convolutional layers total);
2. 512 filters per convolutional layer, for all convolutional layers;
3. size-21 filters in the first convolutional layer;
4. size-75 filters in the deconvolutional layer;
5. counts task weight (lambda) of 100.

### ProCapNet model training

ProCapNet was trained and evaluated using 7-fold cross-validation split by chromosome, with no overlap between training, validation, and test sets. Seven folds were chosen, rather than five, because using additional folds increases the size of the training set for each fold. All downstream analyses, excluding those where models were being evaluated on held-out test sets, used predictions and contribution scores averaged across the 7-fold models.

The set of training examples for the main ProCapNet model consisted of all PRO-cap peaks in the human K562 cell-line as well as background K562 DNase-hypersensitive sites, sampled randomly without replacement from the training set chromosomes. Background DNase-hypersensitive sites were at least 500bp away from the center of any PRO-cap peak and were resampled independently for each training epoch. A 7:1 ratio of PRO-cap peaks to DNase-hypersensitive sites was enforced within batches of size 32. We employed two forms of data augmentation: first, training examples were randomly reverse-complemented with 0.5 probability, and second, each example was centered *±*200bp from the center of the PRO-cap peak (or DNase-hypersensitive site), with the offset selected with uniform probability. All models were trained using early stopping (patience of 10 epochs) which monitored the overall loss on the fold validation set.

For ProCapNet models trained in other cell-lines, the training set consisted of PRO-cap peaks and DNase-hypersensitive sites for that cell-line. For the version of ProCapNet trained on only promoters, the training set only included PRO-cap peaks with a center that fell within 500bp of an ENCODE-annotated candidate promoter. DNase-hypersensitive sites were not used in training Promoter-ProCapNet, as they could include enhancers without high initiation signal.

RAMPAGE-Net was trained identically to ProCapNet, including with 7-fold cross-validation by chromosome and with the same architecture, but to predict RAMPAGE signals in K562 and using RAMPAGE peaks. The training dataset consisted of all RAMPAGE peaks from the training chromosomes plus background regions of DNase peaks that did not overlap RAMPAGE peaks.

ProCapNet was implemented and trained in PyTorch 1.12.1^129^ using the Adam optimizer ^130^ (`learning_rate = 0.0005`).

### Quantifying profile task performance

The Jensen-Shannon Distance (JSD) metric was used to measure how similar each ProCapNet profile task prediction was to the distribution of the measured PRO-cap data. JSD was calculated using the scipy jensenshannon function (base = 2). Because PRO-cap peaks with lower read coverage produce higher (worse) JSD values due to the inherently stochastically sampled nature of sequencing reads, and because model predictions are real-valued numbers and not whole read counts, it was necessary to normalize the JSD metric to remove this skew. We generated two pseudo-replicates for each PRO-cap dataset by splitting the reads randomly into two groups and then used the JSD calculated by comparing the two pseudo-replicates at each PRO-cap peak as the upper bound. We next calculated a lower bound JSD that corresponds to if the model predicted a perfectly flat (uniform) read distribution. Actual measured vs. model-predicted JSD values were then min-max normalized, individually for each example, using these bounds.

### Quantifying initiation directionality and dispersion

The Orientation Index ^83^ (OI), which measures the strand asymmetry of PRO-cap data within a given region, was calculated by dividing the maximum number of reads on either strand by the sum of the reads on both strands. The Normalized Shape Index (NSI), modified from Hoskins *et al*. 2011^84^, measures the dispersion of PRO-cap data within a given region and was calculated as the Shannon entropy of read positions divided by the natural *log*-*log* of the read counts, to reduce skew from differences in coverage. Both metrics were calculated using reads found within a 1,000-bp window centered on each PRO-cap peak.

### cCRE, promoter class, and gene region identification

Annotations of candidate promoters and enhancers in K562 were downloaded from ENCODE (ENCSR301FDP) ^79^. These annotations were curated using orthogonal experimental measurements to PRO-cap, including chromatin accessibility and histone modifications. PRO-cap peaks were labeled as promoters or enhancers according to whether their center was within 500bp of one or more annotated “promoter-like signature” or “distal enhancer-like signature” candidate elements, respectively. Promoter PRO-cap peaks also within 500 bp of “proximal enhancer-like signature” candidate elements were labeled as promoters with proximal enhancers.

For the purposes of stratifying model performance across promoter classes and gene regions, the stratifications were defined as follows. Promoters with TATA boxes were defined as promoter PRO-cap peaks that also contained an identified instance of either of the TATA box motifs identified by TF-MODISCO. RP-TCT promoters were defined as PRO-cap peaks that fell within regions from 800bp upstream to 700bp downstream of the TSSs of all 84 GENCODE-annotated canonical ribosomal protein genes, filtered for close matches to the sequence TCTT.

Housekeeping promoters were defined as in Bergman *et al*. 2022^52^. Protein-coding and lncRNA promoter labels corresponded to GENCODE annotations of gene function.

PRO-cap peaks were stratified into genic, intergenic, intronic, and exonic regions according to cell-type-agnostic GENCODE annotations. First, all peaks that fell within windows upstream 300 bp to downstream 200 bp from any annotated TSS were labeled ”At TSS”. Second, all peaks that were not already labeled “At TSS” were labeled as being in a gene body if their center was located within any GENCODE transcript, or as intergenic otherwise. Third, every peak in a gene body, but not ”At TSS”, was further stratified by whether it fell within any annotated exon (and if so, whether it fell within any annotated UTR); otherwise it was labeled intronic.

For performance stratification across mappability bins, the overall mappability of each PRO-cap peak was defined as the fraction of bases within a 1 kb window centered on the peak that were 100% mappable according to 36-mer multi-read Umap annotation tracks ^126^.

### Contribution scoring and score clustering

Contribution scores were generated for all bases in each sequence in the training and validation set using DeepSHAP ^88^, as implemented in PyTorch Captum v0.5.0. DeepSHAP is an extension of DeepLIFT ^89^ that approximates SHAP values by contrasting the model’s prediction for a given sequence against predictions for a set of “background” reference sequences. We used 25 dinucleotide shuffles of the sequence being scored as the reference sequence set. Because DeepSHAP relies on having a single scalar value to represent model outputs, the output of the profile head was summarized as follows: the logits (profile head output pre-softmax) were mean-normalized per-sequence, and then the dot product between the normalized logits and the post-softmax profile head output was computed. This is equivalent to weighting the logits by the per-base predicted probabilities of TSS positioning, and then summing over the 1,000-bp output window and both strands. For every sequence scored, we took the average of the per-nucleotide, per-base scores (a 4-bases by 2,114-positions sized array) across the forward strand sequence and its reverse complement to improve the robustness of the scores.

To aggregate contribution scores over multiple instances for motif discovery, a performance-improved version of the TF-MODISCO algorithm ^90^, tf-modiscolite v2.0.0, was applied to the DeepSHAP scores from the central 1,000 bp of all scored sequences. The parameters used for tf-modiscolite were:

~~~
max_seqlets_per_metacluster = 1000000,
sliding_window_size = 20,
flank_size = 5,
target_seqlet_fdr = 0.05,
n_leiden_runs = 50.
~~~

TF-MODISCO patterns were matched to known TF motifs by querying the JASPAR database ^131^ of all vertebrate motifs for matches to CWMs using the TOMTOM tool ^132^ from the MEME suite ^133^ and manual curation.

### Motif instance calling

To identify individual instances of the sequence patterns reported by TF-MODISCO, the following procedure was applied: first, all sequence positions within *±*1057 bp of each PRO-cap peak center were scored via convolutional scanning of each motif’s CWM. Second, all positions were scored again by scanning each CWM across the task-specific DeepSHAP contribution scores. Third, we filtered for hits with both high sequence-match scores and high contribution-match scores, with thresholds set manually for each motif by human inspection to ensure proper match fidelity. Overlaps between hits were resolved by choosing the motif instance with the largest contribution-match score multiplied by the motif’s length. Downstream analyses used the set of hits based on the profile task-derived CWMs and contribution scores because they were qualitatively cleaner than CWMs and hits specific to the counts task; hits from the counts task reported in Fig. 2A were derived from counts task contribution scores, but used the profile task-derived CWMs for scanning to allow for fair comparison of motif hit counts across tasks.

### Principal components analysis on model embeddings of promoters and enhancers

Model embeddings were defined as the outputs of the global average pooling layer, which follows the final dilated convolution layer that is shared between the profile and counts task heads. embeddings were generated for all K562 PRO-cap peaks, and principal component analysis was run on this dataset using the PCA module in scikit-learn v1.1.2. PRO-cap peaks annotated as overlapping candidate promoters and distal enhancers were then projected into the PCA space.

### Linear and logistic regression models

All linear models were fit using the ordinary least squares (OLS) function with default parameters from the statsmodels v.0.13.2 Python package. For all linear models, the same 7-fold cross-validation scheme was applied for model fitting vs. evaluation. An intercept term was added during fitting, and the adjusted *R*^2^ on held-out PRO-cap peaks, averaged across folds, was reported.

For linear models fit using chromatin state information, scalar values for accessibility and histone mark signals were calculated per PRO-cap peak by taking the mean of the *log* of fold-change over control values within a 1-kb window centered on the peak center. For models using chromatin state information that were fit with multiple input features (including models using ProCapNet predictions plus one chromatin state signal), all possible second-order interaction terms were included in the model.

Logistic regression models for cCRE category classification were fit using the LogisticRegression module from scikit-learn v1.1.2. Accuracy values reported are averages across the held-out cCRE-overlapping peaks from 7-fold cross-validation.

### Effects of motif ablation on predicted PRO-cap coverage and profile shape

Motif ablation in the analysis of *MYC* promoter epistasis was performed manually: the sequence corresponding to the motif was replaced by a sequence of equal length designed by hand to no longer match that motif, nor create a match to any other ProCapNet-learned motif. All predictions and contribution scores shown are the average of outputs from each of the models trained across 7 folds.

For the systematic analysis of motif ablation across all peaks, the following process was performed at every instance of each motif identified in a PRO-cap peak. First, model predictions were generated for both the original sequence centered at the peak and 25 sequences where the motif instance was replaced with random nucleotides to destroy the motif, with the rest of the sequence remaining intact (**Fig. 5B**). The random nucleotides were sampled with probabilities matching the frequencies of bases in the motif and within *pm*50 bp of flanking sequence around the motif. Second, to summarize the effect of ablation for the counts task, the fold-change was calculated between the unperturbed sequence counts prediction and each of the perturbed sequence counts predictions, and the median fold-change across repeated ablations was reported (**Fig. 5C, bottom**). Third, to summarize the effect of ablation on the profile task, the median model prediction after ablation was computed over all ablations, and then the median was taken across all instances of the motif. Profiles were centered on the motif instances and corrected according to motif orientation (if a motif was in the reverse orientation, the predictions were reversed).

### Software and data availability

Code to download and preprocess all data, train ProCapNet, and reproduce all downstream analyses is available at https://github.com/kundajelab/ProCapNet/.

ProCapNet models, predictions at PRO-cap peaks, contribution scores, TF-MODISCO outputs, and model training files are available through the ENCODE portal, with accession IDs ENCSR740IPL, ENCSR072YCM, ENCSR182QNJ, ENCSR797DEF, ENCSR801ECP, ENCSR860TYZ for K562, A673, Caco-2, Calu3, HUVEC, and MCF10A, respectively. See Table S5 for a complete list of the data available.

## Author contributions

A.K., K.C., and G.K.M. conceived the project. S.R.G, H.Y., and J.T.L. generated the PRO-cap experimental data. M.Y. performed analyses included in Figure 7, and A.M. performed analyses included in Figure 6. K.C. performed all other analyses, with A.K. providing primary conceptual advising and J.S. providing conceptual and implementation advising and code. K.C. primarily drafted the manuscript and figures, with J.S., G.K.M., and A.K. contributing to writing and revisions and J.T.L. providing draft feedback.

## Acknowledgments

We would like to thank all members of the ENCODE consortium for generating and contributing datasets used in this paper. We would like to thank Aman Patel for help with model uploads to the ENCODE portal, and Jennifer Jou, Idan Gabdank and Ben Hitz and other members of the ENCODE DCC for supporting ProCapNet models at the ENCODE portal. We would like to thank Julia Zeitlinger, Jesse Engreitz, William J. Greenleaf, Lacramioara Bintu, and Michael C. Bassik for their insightful feedback on results and project directions.

The authors acknowledge funding support from NIH grants 5U24HG007234, U01HG009431, U01HG012069, and U24HG007234 to A.K. K.C. was supported in part by the Pierre & Christine Lamond Stanford Graduate Fellowship.

## Supplementary Materials

### Supplementary Tables

**Supplementary Table 1:** Overlap of TF-MODISCO patterns resembling transposable elements with annotated repeats.

| TE Pattern | Repeat Overlap | Most Common Repeat Family | Frac. Repeat Overlap Matching Family |
| --- | --- | --- | --- |
| Profile TE Pattern 1 | 82% | ERV1 (LTR) | 100% |
| Profile TE Pattern 2 | 84% | ERVL-MaLR (LTR) | 97% |
| Profile TE Pattern 3 | 81% | ERV1 (LTR) | 100% |
| Profile TE Pattern 4 | 91% | ERV1 (LTR) | 99% |
| Profile TE Pattern 5 | 75% | Alu (SINE) | 93% |
| Profile TE Pattern 6 | 81% | ERVL-MaLR (LTR) | 100% |
| Profile TE Pattern 7 | 70% | ERV1 (LTR) | 100% |
| Profile TE Pattern 8 | 78% | Alu (SINE) | 94% |
| Profile TE Pattern 9 | 67% | ERV1 (LTR) | 93% |
| Counts TE Pattern 1 | 96% | ERV1 (LTR) | 100% |
| Counts TE Pattern 2 | 82% | L1 (LINE) | 97% |
| Counts TE Pattern 3 | 82% | ERVL-MaLR (LTR) | 98% |
| Counts TE Pattern 4 | 73% | L1 (LINE) | 96% |
| Counts TE Pattern 5 | 95% | ERVL-MaLR (LTR) | 96% |
| Counts TE Pattern 6 | 91% | Alu (SINE) | 97% |
| Counts TE Pattern 7 | 85% | ERV1 (LTR) | 86% |
| Counts TE Pattern 8 | 73% | ERVL-MaLR (LTR) | 90% |
| Counts TE Pattern 9 | 82% | ERV1 (LTR) | 93% |
| Counts TE Pattern 10 | 61% | Alu (SINE) | 94% |

**Supplementary Table 2:** Specificity of motif instances identified using contribution scores of the ProCapNet profile task.

| Motif | Hits From Sequence Scanning | Hits After Score Filtering | Fraction of Sequence Hits Retained |
| --- | --- | --- | --- |
| SP1/BRE | 451541 | 92744 | 0.21 |
| CA-Inr | 8304122 | 56605 | 0.01 |
| ETS | 86478 | 23814 | 0.28 |
| NFY | 145277 | 19495 | 0.13 |
| NRF1 | 124318 | 36430 | 0.29 |
| ATF1 | 64902 | 5789 | 0.09 |
| TATA | 126943 | 7436 | 0.06 |
| THAP11 | 6224 | 2318 | 0.37 |
| YY1 | 2586 | 1398 | 0.54 |
| AP1 | 61128 | 11630 | 0.19 |
| TA-Inr | 4812930 | 25569 | 0.01 |
| CTCF | 2109 | 1019 | 0.48 |
| ZBTB33 | 8202 | 2536 | 0.31 |
| TCT | 701749 | 423 | < 0.01 |
| TATATA | 299303 | 14659 | 0.05 |

**Supplementary Table 3:** Comparison of model predictions and contribution scores between replicate ProCapNets and Promoter-ProCapNet. Values in parentheses indicate standard deviations across folds.

| Models Compared<br>Mean Correlation | Replicate ProCapNets |  | ProCapNet vs. Promoter-ProCapNet |  |
| --- | --- | --- | --- | --- |
| | Pearson $r$ | Spearman $r$ | Pearson $r$ | Spearman $r$ |
| Counts Predictions | 0.976 (0.005) | 0.974 (0.005) | 0.916 (0.015) | 0.941 (0.011) |
| Profile Predictions | 0.898 (0.004) | 0.943 (0.002) | 0.881 (0.006) | 0.930 (0.006) |
| Counts Contribution Scores | 0.844 (0.018) | 0.807 (0.020) | 0.692 (0.033) | 0.659 (0.030) |
| Profile Contribution Scores | 0.875 (0.005) | 0.775 (0.007) | 0.840 (0.010) | 0.726 (0.021) |

**Supplementary Table 4:** Additional datasets used in chromatin state analysis for the K562, A673, Caco-2, and HUVEC cell types.

| Cell Type | K562 | A673 | Caco-2 | HUVEC |
| --- | --- | --- | --- | --- |
| DNase-seq | ENCSR000EKS | ENCSR114QAK | ENCSR255STJ | ENCSR000EOQ |
| H3K27me3 | ENCSR000AKQ | ENCSR747BYL | ENCSR000DQL | ENCSR000AKK |
| H3K4me1 | ENCSR000AKS | ENCSR521IZK | ENCSR061UOM | ENCSR000AKL |
| H3K27ac | ENCSR000AKP | ENCSR714TJD | NA | ENCSR000ALB |
| H3K9me3 | ENCSR000APE | ENCSR988EGR | ENCSR401WUV | ENCSR000ATB |
| H3K36me3 | ENCSR000AKR | ENCSR581PUR | ENCSR000DQK | ENCSR000ALC |
| H3K4me3 | ENCSR668LDD | ENCSR435FGK | ENCSR000DQM | ENCSR000AKN |

**Supplementary Table 5:** ENC-IDs for ProCapNet data available through the ENCODE portal.

| Data | K562 | A673 | Caco-2 | Calu3 | HUVEC | MCF10A |
| --- | --- | --- | --- | --- | --- | --- |
| all data | SR740IPL | SR072YCM | SR182QNJ | SR797DEF | SR801ECP | SR860TYZ |
| training and test regions tar | FF033LJA | FF611RQM | FF603POZ | FF571XHE | FF065FTI | FF941RDG |
| models tar | FF976FHE | FF508EUP | FF575XTL | FF180BYW | FF822XKO | FF568EGA |
| observed signal (+ strand) bigWig | FF798GNW | FF484JMU | FF290ILU | FF726ZFO | FF515GWR | FF460DXE |
| observed signal (- strand) bigWig | FF662SHP | FF248NRG | FF674WYH | FF694ACJ | FF932EJP | FF466PYB |
| predicted signal (+ strand) bigWig | FF810TSX | FF245QOS | FF781NNR | FF297DYW | FF522DPN | FF413SKO |
| predicted signal (- strand) bigWig | FF977ZRF | FF568EMG | FF512NRZ | FF726MZN | FF130FIR | FF751HYG |
| profile contribution scores bigWig | FF105JTF | FF395XCX | FF566GOV | FF369BLK | FF090ECF | FF117LHD |
| profile contribution scores tar | FF407PRC | FF993LJR | FF925QDB | FF530ZUR | FF846XLY | FF282QWJ |
| counts contribution scores bigWig | FF399XSL | FF496KXS | FF773CZW | FF255ZJK | FF142HSZ | FF898JHY |
| counts contribution scores tar | FF186EVD | FF205ATX | FF674PEC | FF838EMK | FF956VWD | FF220LKE |
| predicted + scored regions bed | FF271LOH | FF605QYF | FF594CPW | FF420WIU | FF145VXT | FF187MYU |
| motifs tar | FF804ZPG | FF630EMW | FF982OZL | FF083YAU | FF276YTU | FF589NBY |
| motif instances bed | FF070AVM | FF358WWO | FF935ACF | FF075CBM | FF139KSH | FF693EHJ |
| motif instances bigBed | FF624OCH | FF890XJG | FF214LNU | FF351WJS | FF479KIM | FF264QWB |
| motifs report tar | FF608IHS | FF710OHX | FF702DZH | FF832NQX | FF316GZQ | FF145DCV |

### Supplementary Figures

**Supplementary Figure 1:**
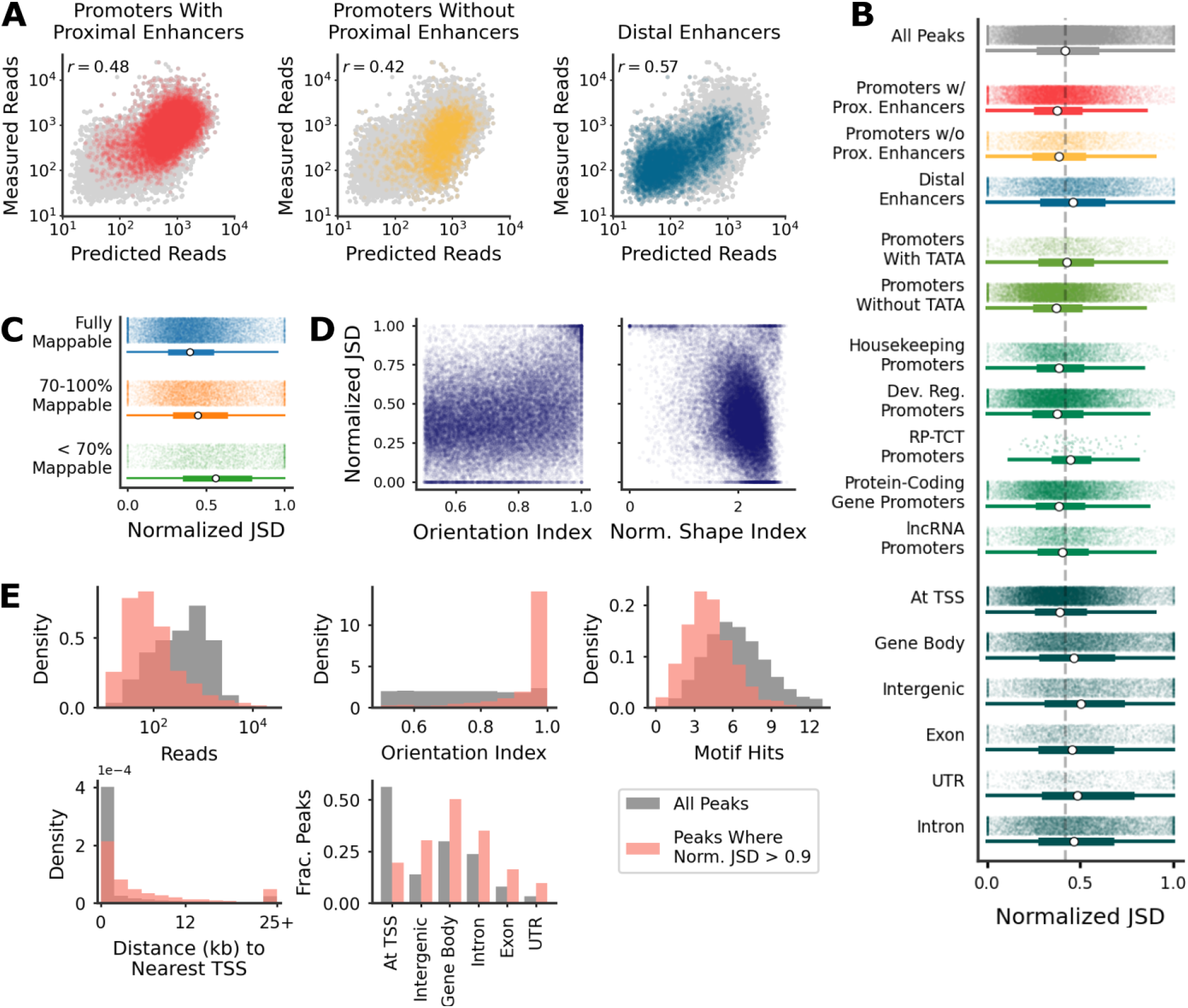
Model performance across stratifications of PRO-cap peaks. (A) Predicted and measured overall PRO-cap signal on held-out test peaks across all model folds, colored by various categories of CRE (*cis*-regulatory element). (B) ProCapNet profile task performance distributions across various subsets of PRO-cap peaks. Dev. Reg., developmentally regulated (non-housekeeping); RP-TCT, ribosomal protein gene promoters with TCT-like sequences. Dashed line indicates the median for all peaks; white dots indicate individual group medians. (C) ProCapNet profile task performance distributions for fully mappable (100% umap track coverage of 1), mostly mappable (umap track coverage of 1 between 70 and 100%), and least mappable (umap track coverage of 1 less than 70%) PRO-cap peaks. (D) Profile task performance as a function of strand asymmetry (Orientation Index) and dispersion of TSSs (Normalized Shape Index; see Methods for detailed definitions). (E) Statistics across all PRO-cap peaks vs. the subset of PRO-cap peaks where ProCapNet profile task performance, measured by normalized JSD, was greater than 0.9.

**Supplementary Figure 2:**
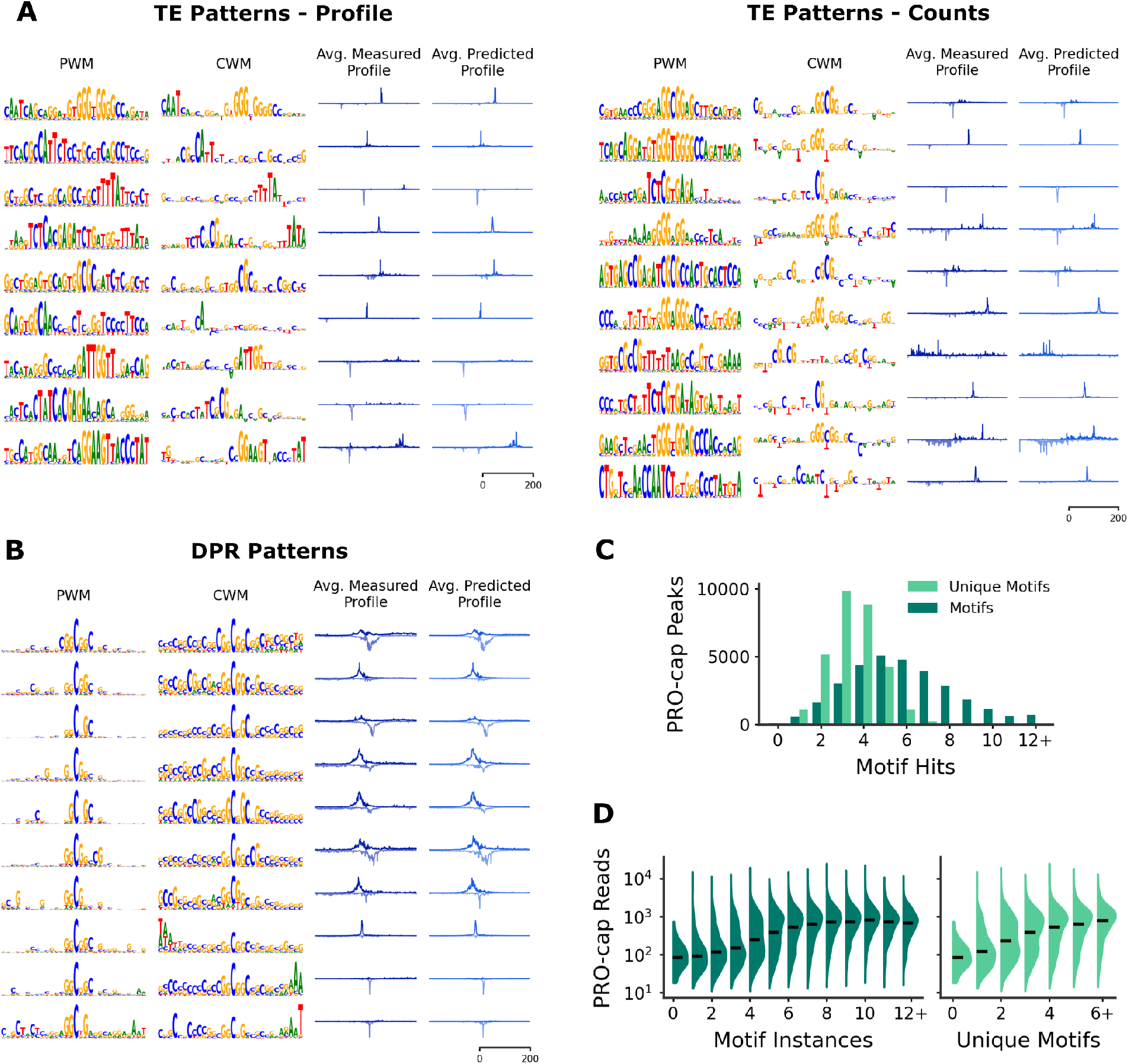
Additional TF-MODISCO results and motif complexity in PRO-cap peaks. (A) Patterns resembling putative transposable elements (TEs) found by TF-MODISCO for the ProCapNet profile task (left) and counts task (right). (B) GC-rich patterns found by TF-MODISCO in the downstream promoter region (DPR) that were highlighted by high contribution scores for the ProCapNet profile task.(C) Histogram of motif instances identified in PRO-cap peaks, either including or not including homotypic motif multiplicity. (D) Distribution of PRO-cap read counts at peaks with increasing motif instances. Black lines indicate median values.

**Supplementary Figure 3:**
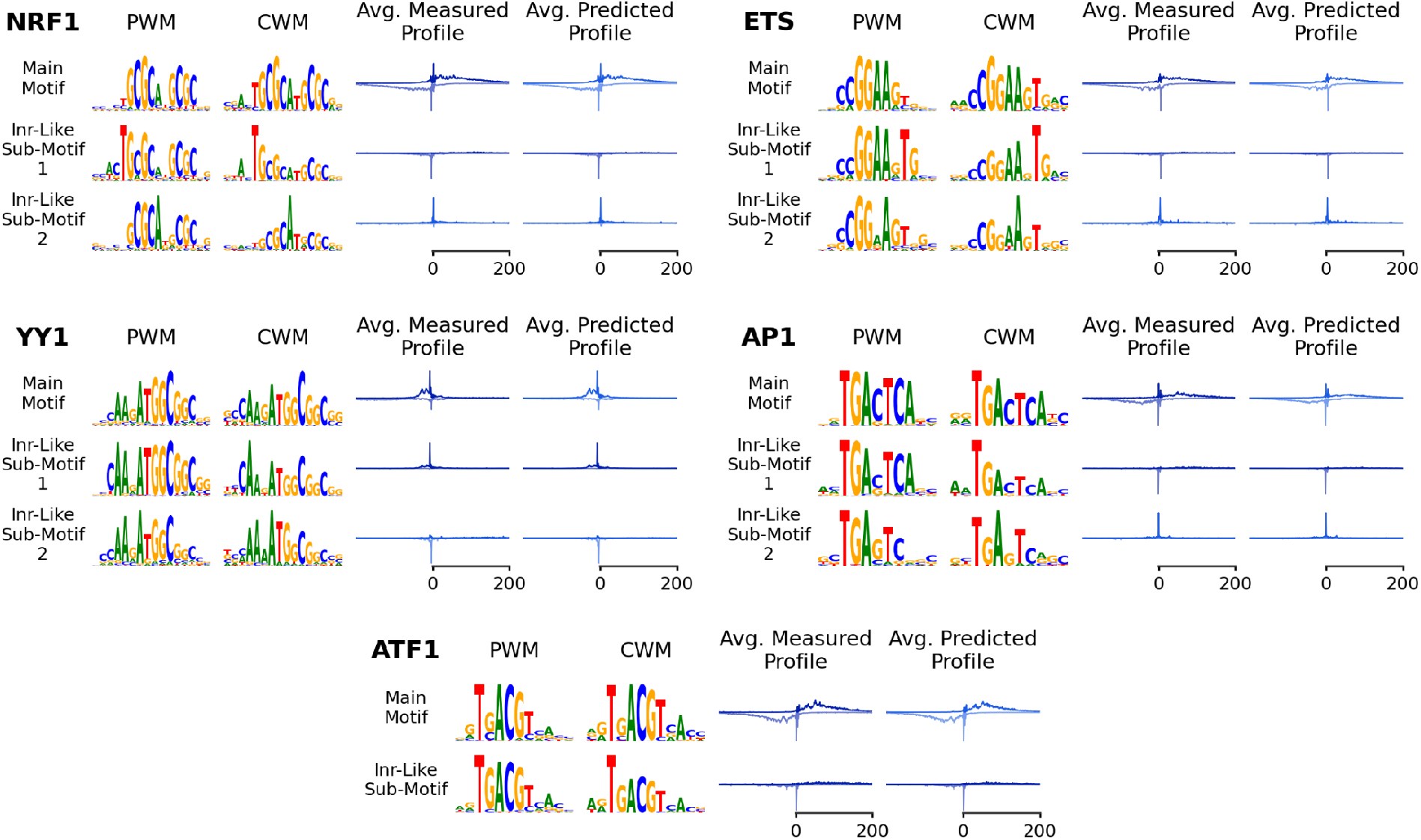
Inr-like TSS positioning secondary functions identified by TF-MODISCO subpatterns for several motifs. PWM/CWM representations and average profiles are shown for 1) the set of highest-scoring motif instances used in subclustering by TF-MODISCO, and 2) each subcluster derived from those highest-scoring instances where an Inr-like submotif was emphasized in the CWM and the average profile indicated direct initiation activity.

**Supplementary Figure 4:**
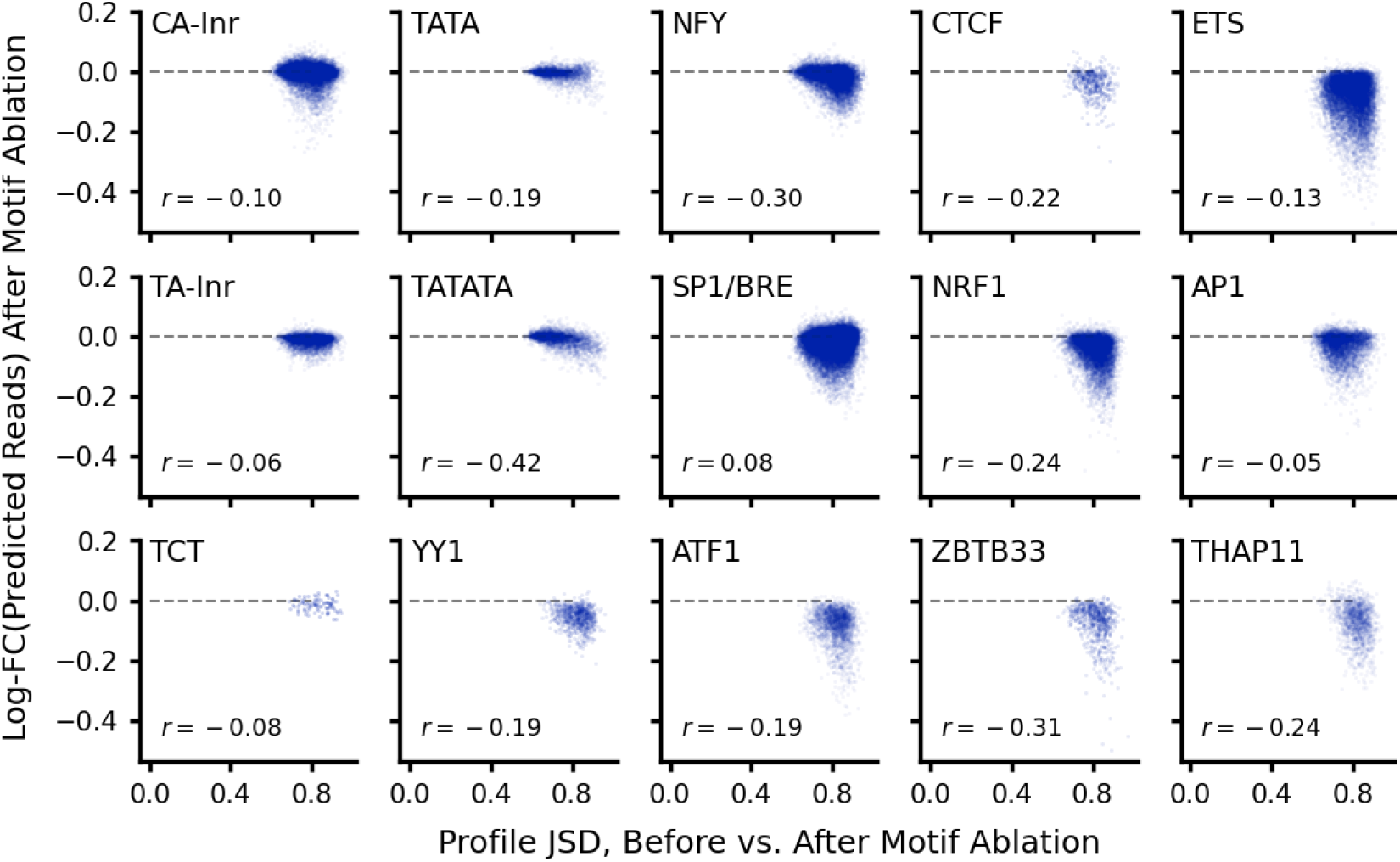
Predicted effects of motif ablation on TSS positioning (x-axis) vs. overall initiation activity (y-axis). *r*, Spearman correlation. Dashed line indicates a fold-change of 1.

**Supplementary Figure 5:**
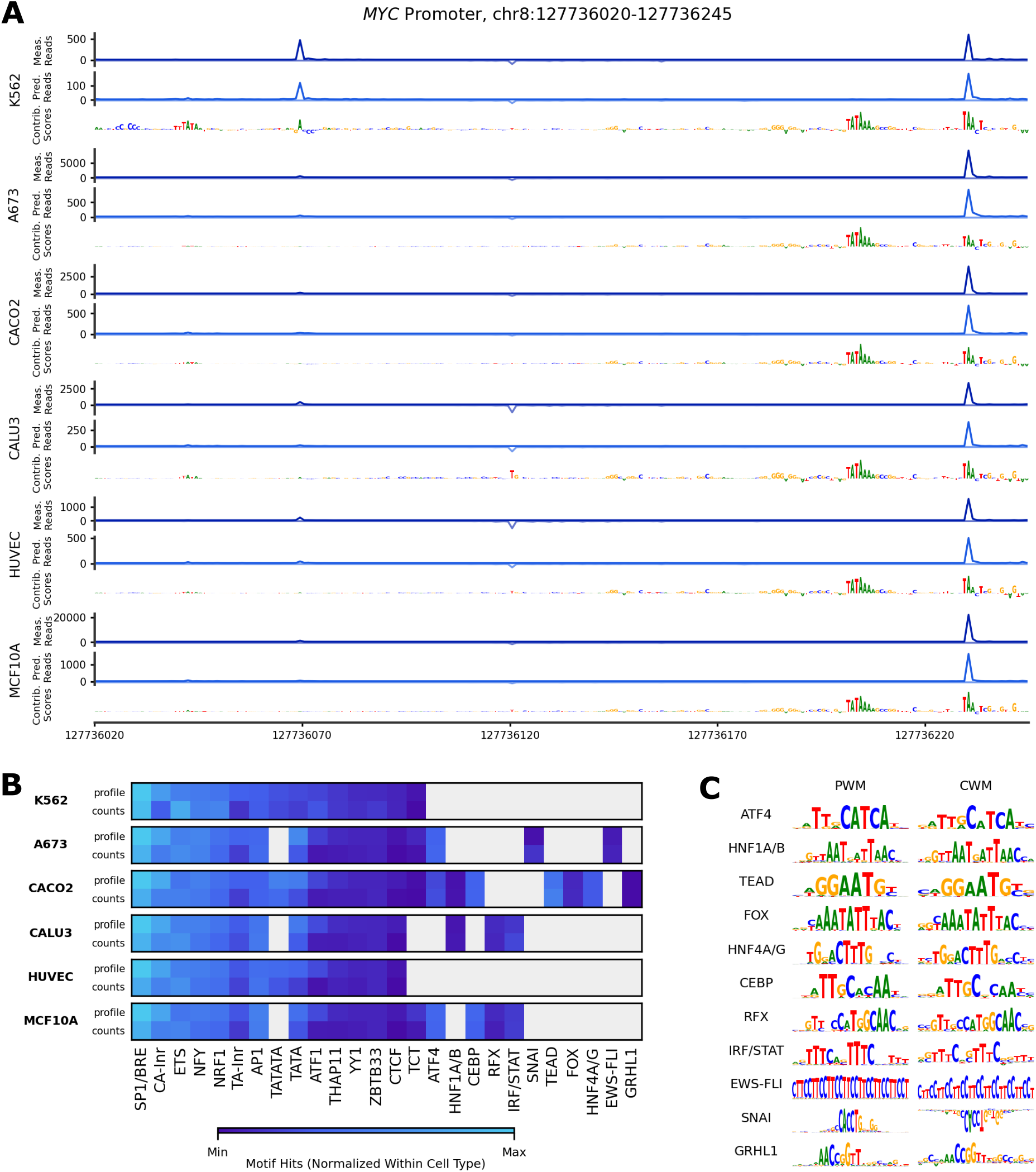
Example predictions and discovered motifs from models trained on all available cell types. (A) Measured PRO-cap and ProCapNet predictions and profile task contribution scores at the *MYC* promoter, across all cell types. (B) Motif hit counts across all cell types. (C) Position-weight matrix (PWM) and contribution-weight matrix (CWM) representations of novel motifs discovered in cell types beyond K562. The SNAI CWM’s orientation indicates negative contribution scores, suggestive of repression.

## Supplementary Note: Comparison to the Puffin Model

The Puffin model ^1^, like ProCapNet, aims to predict base-resolution initiation profiles from local DNA sequence, although the two models take different approaches toward achieving this goal.

Puffin uses an innovative strategy to build and train a highly constrained and transparent neural network architecture with directly interpretable parameters ^1^. Puffin’s design enables one to “read-off” many interesting properties of the initiation code from the model parameters and activations. However, there are several ad-hoc aspects to how the Puffin architecture is handcrafted and trained. The model is designed to learn motifs in its first convolution layer such that the identity and length of motifs is explicitly constrained based on a set of motifs curated from the larger, long-context, black-box Puffin-D model. These constraints are iteratively refined via multiple rounds of training and ad-hoc curation. Some motifs are given special consideration without clear justification. For example, Inrs and tri-nucleotide patterns are incorporated in later layers compared to other motifs. Baked into the Puffin architecture is a strong assumption that motif instances impact initiation independently and additively in a position-specific manner. While this strategy appears to converge to models that learn reproducible features, it is unclear how these specific choices of constraints, relative to equally plausible alternatives, affect performance and other downstream conclusions. Also, it is unclear if these design choices are in fact necessary for enhancing discovery of the *cis*-regulatory code of initiation.

In contrast, ProCapNet is based on the “black-box” BPNet neural network architecture, which places no explicit constraints on the number or properties of motifs and higher-order syntax that the model can learn. Our model architecture is “black-box” in terms of model parameters not being directly interpretable. However, we show that well-established post-hoc model interpretation methods such as DeepLIFT, TF-MODISCO and *in silico* counterfactual perturbation experiments provide deep insights into the *cis*-regulatory code of initiation learned by the model. Further, model training and post-hoc interpretation approaches do not require manual intervention or ad-hoc curation, making ProCapNet’s strategy inherently more replicable and scalable. For example, ProCapNet can easily be trained on PRO-cap data from diverse cell types without any manual intervention, since the model can automatically adapt to learning cell-type-specific features and syntax as needed. Puffin, on the other hand, would require separate, semi-manual motif curation and architecture tweaks for each cell type. Our approach is based on the philosophy that constraining architectures based on incomplete prior knowledge carries the risk of overriding the model’s ability to learn novel sequence features and syntax, thereby potentially hindering a faithful explanation of its predictions. This effectiveness of this paradigm is strongly supported by several prior applications of BPNet and its derivatives to TF binding and chromatin accessibility profiles ^2,3^.

Here, we present a direct comparison of Puffin to ProCapNet to understand pros and cons of each approach. We discuss these and other differences between ProCapNet and Puffin and their consequences on predictive performance and interpretation in more detail below. We specifically restrict our comparisons to the Puffin PRO-cap model, rather than the larger Puffin-D model, since (1) Puffin is contextually equivalent to ProCapNet, and (2) the Puffin study exclusively uses the Puffin model for all downstream analyses of the *cis*-regulatory code of initiation.

### ProCapNet slightly outperforms Puffin at predicting initiation profiles from Puffin’s cell-type-agnostic, aggregated dataset

To assess if Puffin’s design choices result in any predictive performance advantage over ProCapNet, we re-trained ProCapNet on Puffin’s dataset, which consists of cell-type-agnostic profiles averaged over a large collection of cell types. We used the exact set of genomic regions used to train Puffin, and then evaluated both models on Puffin’s test set. We chose to re-train ProCapNet on the Puffin dataset, rather than re-training Puffin on cell-type-specific data, due to the difficulties of replicating Puffin’s complex, iterative model design and training approach, discussed above. ProCapNet slightly outperformed Puffin, with average JSDs of 0.58 vs. 0.61 (lower is better), respectively (**Fig. S6A**). Thus, Puffin does not provide any prediction performance advantage over ProCapNet.

### ProCapNet significantly outperforms Puffin at predicting and interpreting cell-type-specific TSSs

Unlike Puffin, which is trained on merged PRO-cap profiles aggregated across diverse cell contexts, ProCapNet models are trained specifically for each cell context. This approach enables ProCapNet to capture both shared and unique motifs, including those not accounted for by Puffin, such as CTCF, certain Inr variants, and all the other cell-type-specific ProCapNet motifs (**Fig. S5B**).

Since ProCapNet (like Puffin) only models local sequence context, it cannot capture the cell-type-specific influence of distal regulatory elements, resulting in largely cell-type-invariant predictions of initiation activity (**Fig. 7**). How ever, ProCapNet is still able to predict cell-type-specific positioning of TSSs, which Puffin inherently cannot (**Fig. S5A**).

**Supplementary Figure 6:**
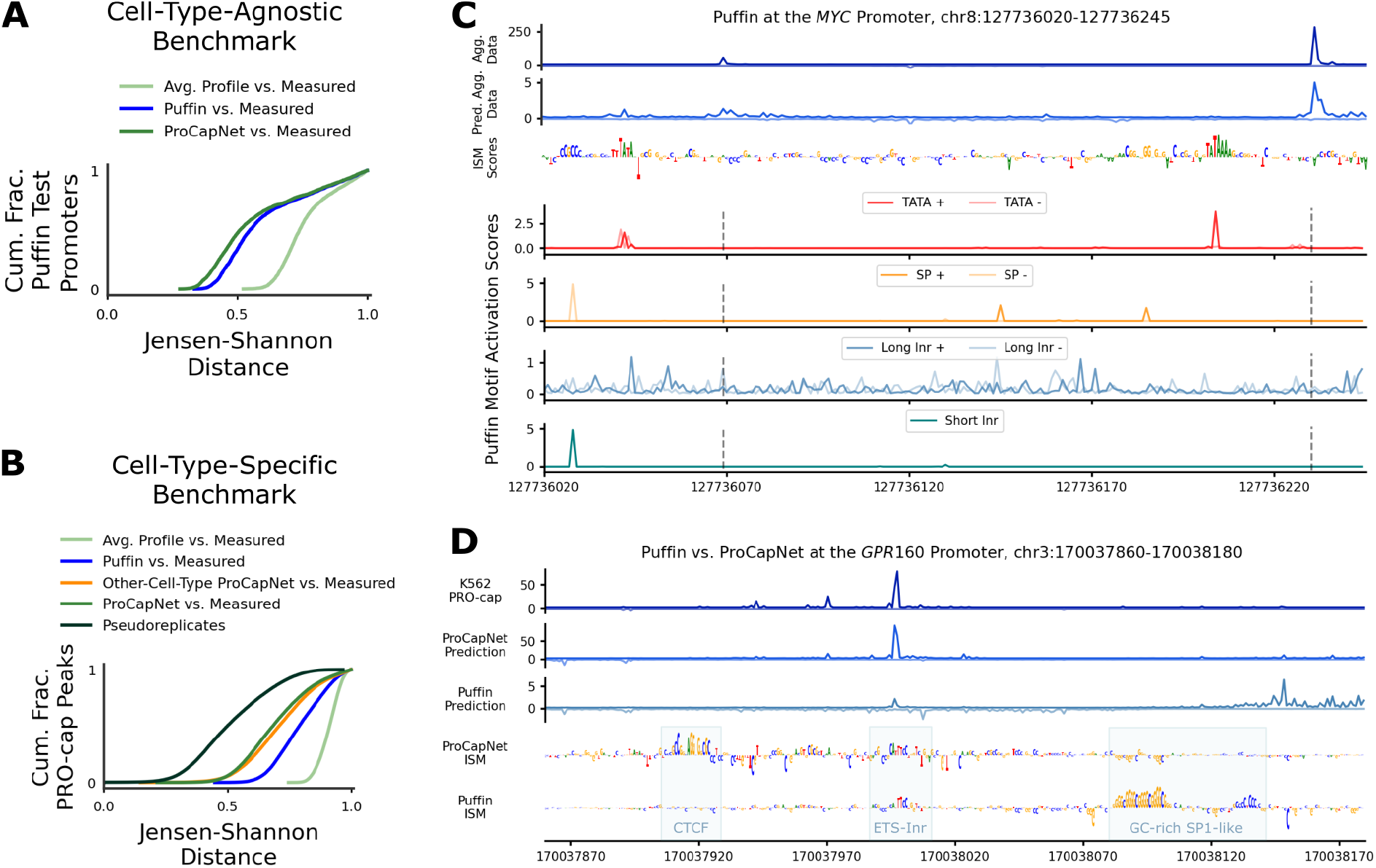
Puffin vs. ProCapNet TSS positioning predictive performance benchmarks, and Puffin interpretation at the *MYC* locus. (A) Predictive performance on Puffin’s test set for Puffin vs. a version of ProCapNet re-trained on Puffin’s training data. (B) Predictive performance for Puffin vs. the K562-trained ProCapNet on K562 PRO-cap across all peaks (using cross-validation for ProCapNet). “Other-Cell-Type ProCapNet” refers to the A673-trained ProCapNet. (C) Puffin’s aggregate, cell-type-merged data (“Agg. Data”), Puffin predictions, Puffin ISM contribution scores, and Puffin-derived motif activation scores for relevant motifs at the *MYC* promoter. Dashed vertical lines indicate the two PRO-cap sense TSSs. (D) Measured K562 PRO-cap, model predictions, and ISM contribution scores from Puffin and ProCapNet at the *GPR160* promoter.

We systematically compared the performance of Puffin’s cell-type-agnostic model to cell-type-specific ProCapNet models at predicting cell-type-specific PRO-cap profiles. Initial benchmarks against K562 PRO-cap profiles demonstrated that ProCapNet models trained on K562 profiles substantially outperformed Puffin, highlighting the importance of cell-type-specific training (**Fig. S6B**). These evaluations were performed across all K562 PRO-cap peaks using the original cross-validation scheme used for ProCapNet. This set up causes some inadvertent train-test leakage in favor of Puffin, since some of the peak sequences in ProCapNet’s test sets are present in Puffin’s original training set, although Puffin is trained on aggregated profiles. Even when ProCapNet was trained on disparate cell-line data (A673, which is a sarcoma cell-line), it still surpassed Puffin at predicting K562 PRO-cap profiles, indicating superior transferability of cell-type-specific models over aggregated models (**Fig. S6B**). Aggregating profiles across cell types may introduce unexpected artefacts, resulting in predictions that are out-of-distribution with respect to profiles from any cell type.

These issues with prediction of cell-type-specific profiles introduce further downstream challenges with interpretation. For example, at the *MYC* promoter, the upstream TSS is specifically active in K562 but not in all the other ENCODE cell-lines with PRO-cap data. ProCapNet models from various cell types accurately predict the cell-type-specific activity of this TSS, with sequence contribution scores derived from each model reflecting the differential influence of motifs around this TSS (**Fig. S5A**). In contrast, the K562-specific upstream TSS shows weak signal in the aggregated profiles used to train Puffin and in Puffin’s predictions (**Fig. S6C**). Aggregation thus obscures the true distribution of activity of TSSs across cell types, especially for strong but cell-type-specific TSSs, making it unclear whether a TSS’s weak aggregate activity reflects uniform low activity across all cell types or high activity in only a few. Similarly, interpreting sequence features derived from cell-type-agnostic models introduces ambiguity. For instance, if a motif contributes to initiation at the upstream TSS with weak aggregate activity, it’s unclear whether the motif generally has a weak effect or if its impact is cell-type-specific. Only models trained on cell-type-specific datasets can circumvent this issue, providing outputs and interpretations that accurately reflect the transcriptional state and its underlying sequence drivers in specific cell contexts.

### Motif epistasis learned by ProCapNet improves prediction and interpretation of initiation over Puffin’s additive model

Puffin employs a transparent model architecture where motifs are forced to contribute independently and additively to TSS positioning. ProCapNet’s architecture does not impose explicit additive constraints, giving the model the flexibility to learn epistatic interactions if needed. Our post-hoc model interpretation methods show that ProCapNet indeed learns complex epistatic interactions among motifs and TSSs, which appear to be crucial for accurate prediction of transcription initiation activity and TSS positioning (**Fig. 4, Fig. 5**). Our results contradict the assumptions of additivity and independence encoded in the Puffin model. Experimental evidence supports the significance of epistatic interactions among motifs in transcription initiation, showing that a motif’s position relative to the TSS, its orientation, and the surrounding sequence context significantly influence its regulatory impact ^4^.

We present two case studies that support the key role of motif epistasis missed by Puffin.

Initiator (CA-Inr, TA-Inr) elements play a well-known role in positioning TSSs. These are very short motifs, and due to this, strong matches to their consensus sequence are ubiquitous across the genome. However, only a tiny fraction of these instances are associated with transcription. ProCapNet clearly identifies Inr elements co-localized with a large fraction of highly transcribed TSSs (**Fig. 2**). We also find that the Inr is exceptionally epistatic and that interactions with other motifs in the proximal sequence context strongly affect the influence of each Inr on positioning (**Fig. 4**). Puffin cannot model such interactions, which leads it to frequently miss active Inrs at well-positioned, strongly active TSSs that are clearly identified by ProCapNet. This issue occurs despite Puffin explicitly encoding both a short and a long version of the Inr in the model. This limitation becomes apparent in our comparative analyses of the *MYC* promoter, where Puffin fails to detect multiple Inr elements identified by ProCapNet which co-localize with PRO-cap TSSs (**Fig. 4** for ProCapNet, **Fig. S6C** for Puffin). Puffin’s activation scores for the long and short Inr motifs across this locus did not highlight any of the active Inrs found by ProCapNet (**Fig. S6C**, bottom two tracks). We also applied *in-silico* mutagenesis (ISM) for model interpretation across the promoter sequence and further confirmed that Puffin did not predict any effects of mutations at the Inr motif instances (**Fig. S6C**, third track). However, Puffin was sensitive to mutations in the TATA box and SP1/BRE motifs; hence, the lack of sensitivity is very specific to the Inrs. In contrast, ProCapNet clearly identifies the TATA box and SP1/BRE motifs and shows strong epistatic interactions between tham and the Inrs. Since Puffin cannot learn these interactions, it cannot specifically highlight these active Inrs. This lack of detection could also be exacerbated due to these Inr instances not matching the canonical Inr motif CANT; the upstream Inr has only the +1 A, while the downstream Inr matches the TA variant only identified by ProCapNet.

The *GPR160* promoter is another case study showcasing the importance of motif epistasis beyond Inrs. At this locus, ProCapNet and Puffin models significantly differ in their predictions, with ProCapNet more closely mirroring the measured PRO-cap profiles (**Fig. S6D**). We applied ISM, a model-agnostic approach, to identify salient sequence features driving both Puffin and ProCapNet predictions. This analysis highlighted distinct interpretations by the two models, underscoring ProCapNet’s ability to recognize complex, epistatic interactions:

1. **CTCF motif** : ProCapNet’s ISM scores prominently identify a CTCF motif upstream of the TSS, indicative of its role in transcription regulation at this site. Conversely, Puffin is insensitive to the CTCF motif, reflecting its absence from Puffin’s predefined motif set, which underscores the limitation of not allowing the model to autonomously learn and adapt motifs.
2. **SP1/BRE motifs**: Puffin incorrectly predicts substantial initiation activity downstream of the observed TSS, localized near its high ISM scores for multiple GC-rich sequences matching the SP1/BRE motif consensus, which would inadvertently drive activity in a purely additive model. ProCapNet assigns no contribution to these GC-rich sequences, likely due to their position (downstream of the primary TSS-predictive sequence region), a logic that would be encoded via context-aware epistatic interactions in the model.
3. **Dual function Inr-ETS motif** : ProCapNet predicts that the main TSS in this locus co-localizes with with a cryptic CA-Inr element that is intertwined within an ETS motif. ProCapNet is able to learn these dual-role, context-specific cryptic initiators because it has the flexibility to learn motifs and context-aware epistatic interactions (**Fig. S3**). In contrast, Puffin’s constraints limit its ability to detect these cryptic initiators, thereby missing their strong contribution to positioning initiation at this TSS.

These examples illustrate the advantages of ProCapNet’s flexible “blackbox” architecture coupled with post-hoc interpretation methods to accurately predict and interpret transcription initiation, especially in scenarios involving complex motif interactions.

### ProCapNet enables interpretation of sequence drivers of initiation rates and TSS positioning

Finally, ProCapNet models both the overall initiation activity over each sequence and the precise, base-resolution positioning of TSSs captured by the profile shape. In contrast, Puffin claims to only predict profile shape and restricts all downstream analyses to sequence drivers of profile shape, even though it includes activity prediction in its loss function. Using ProCapNet, we explicitly analyze the contribution of sequence features to activity and profile shape, thereby clearly revealing preferential effects of some motifs and higher-order syntax on activation and others on positioning (**Fig. 2, Fig. 5**). The ability to predict overall initiation activity of a sequence is also crucial for design of synthetic promoters with specific transcription output levels and quantification of promoter strength more generally. In conclusion, although Puffin’s innovative model design enables transparent interpretation, the constraints hard-wired into its design, coupled with its training on aggregated, cell-type-agnostic profiles, lead to several shortcomings in predicting and understanding the *cis*-regulatory code of initiation. ProCapNet’s flexible architecture and interpretation framework offers several advantages, highlighting a key point that transparent models does not necessarily yield deeper biological insights.

